# A specialized SUV3 RNA helicase is involved in plant mitochondrial DNA repair

**DOI:** 10.64898/2026.10.23.761044

**Authors:** Moira G. Rodriguez, Claudia D. Raygoza, Ana M. Martinez-Hottovy, David M. Campbell, Emma R. Purfeerst, Luis G. Brieba, Alan C. Christensen

**Author notes:** Moira G. Rodriguez and Claudia D. Raygoza contributed equally to this work.

## Abstract

Synonymous substitution rates in the coding regions of land plant mitochondrial genomes are substantially lower than those in the nucleus. In contrast, non-coding regions are difficult or impossible to align except among very closely related species, suggesting that point mutation rates in these regions are considerably higher. Rearrangement breakpoints occur frequently within non-coding DNA, generating duplications, deletions, and inversions. To reconcile these contradictory patterns of genome evolution, it has been proposed that DNA damage is converted into double-strand breaks that are repaired through a combination of template-directed mechanisms and non-homologous end joining, while purifying selection preserves the integrity of coding sequences. Here we characterized *AtSUV3*, a mitochondrially targeted RNA helicase encoded by *Arabidopsis thaliana At4g14790,* a gene previously proposed to function as part of the plant mitochondrial degradosome. Mutations in *AtSUV3* confer hypersensitivity to ciprofloxacin-induced mitochondrial DNA damage, CRISPR-knockout lines display increased ectopic recombination in the mitochondrial genome, and recombinant AtSUV3 exhibits ATP-dependent RNA helicase activity with a preference for 3′-ended substrates. Together, these findings identify AtSUV3 as a factor involved in mitochondrial dsDNA break repair. Consistent with our results, Arabidopsis and other plants encode an uncharacterized gene, AT5G39840.1, with 75% amino acid similarity with AtSUV3, dubbed herein as SUV3-like. We hypothesize that the AtSUV3 paralog has undergone specialization towards DNA repair, whereas AtSUV3-like protein may play a role in the plant organellar degradosome. Collectively, these findings identify AtSUV3 as a previously unrecognized factor involved in mitochondrial dsDNA break repair and provide a framework for understanding the diversification of DNA repair proteins during plant evolution.

**GRAPHICAL ABSTRACT:** 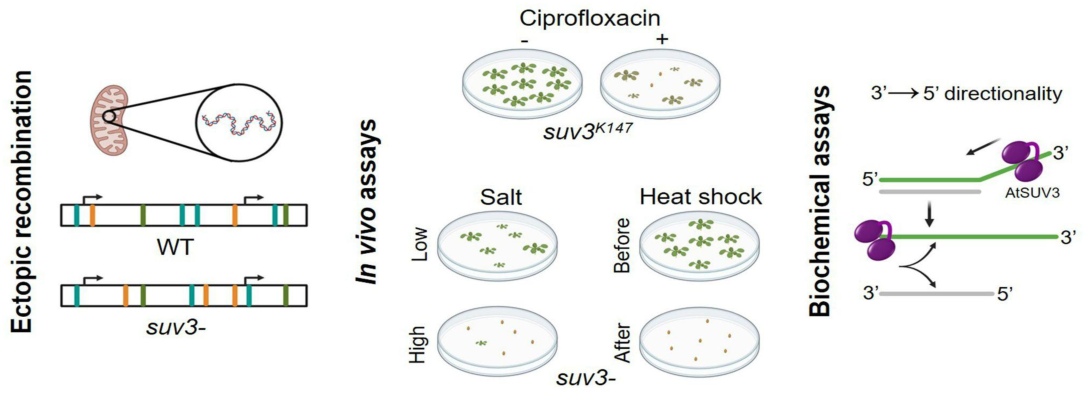

## INTRODUCTION

How do plant mitochondria maintain their genomes so accurately? This has been a puzzle since Palmer and Herbon (1) showed the paradoxical result that mutation rates in genes are very low while rearrangement rates are high (2). Measurements of synonymous substitution rates, which serve as a proxy for the neutral mutation rate, indicate that plant mitochondrial genomes accumulate mutations at a lower rate than the nuclear genome (3). Adding to this puzzle, plant organellar DNA polymerases exhibit lower replication fidelity than the human mitochondrial DNA polymerase POLG, whose errors are associated with high mutation rates during aging (4,5).

In plant mitochondria, genomic rearrangements are common enough that synteny is largely lost between different groups of plants. Rearrangement breakpoints are typically in non-coding sequences, unless a partial or complete duplication of the coding sequence occurs, this is presumably due to selection against breakpoints that disrupt an essential gene function. These partial duplications of genes or small regions of the genome then become non-tandem repeats of various sizes, with most angiosperm genomes having at least dozens of non-tandem repeats of 100-1000 bp (6). Large repeats are less abundant, but many plant mitochondria have at least one pair of non-tandem repeats larger than 1 kb, and these invariably participate in homologous recombination and isomerize the genome (7,8). The 100-1000bp non-tandem dispersed repeats only occasionally recombine in wild type plants, however their recombination can be detected by Southern blotting or PCR (9–11) and are understood to be an indication of a defect in DNA repair pathways (Wallet, Le Ret et al. 2015, Gualberto and Newton 2017, Schatz-Daas, Fertet et al. 2022) or the induction of mitochondrial DNA damage (12–19).

It has been suggested that double-strand break (DSB) repair may be largely responsible for DNA maintenance in plant mitochondria (20). The hypothesis is that nearly all DNA damage is converted to DSBs (possibly by the novel plant-specific MSH1 protein) which are then repaired either by inaccurate end joining or very accurate template-mediated gene conversion or homologous recombination (21–23). Plant mitochondrial genomes are present in many copies per cell (24–26) providing abundant potential repair templates. Selection is expected to eliminate inaccurate repairs that disrupt gene function, while tolerating rearrangements that do not impair gene function. However, many components of the mitochondrial double-strand break (DSB) repair machinery remain unidentified. A recent review of DNA repair in plant mitochondria noted the absence of identified sliding clamps, clamp loaders, endonucleases, and Holliday junction resolvases, while only two DNA helicases homologous to bacterial RadA and RecG associated with DSB repair have been characterized (27). Thus, a more complete understanding of DSB repair, a process central to mitochondrial genome structure and evolution in plants, will require the identification of additional pathway components.

One hallmark of this system is its sensitivity to mutations in nuclear-encoded DNA repair genes, as the mitochondrial genome encodes no known DNA maintenance proteins, allowing aberrant or unrepaired genomes to accumulate. This system is also sensitive to DNA-damaging agents. Ciprofloxacin is an antibiotic that inhibits the organellar DNA gyrase in plants generating DSB via replisome stalling (28), and *Arabidopsis* plants with mutations in known mitochondrial repair genes are hypersensitive to ciprofloxacin (CIP) (13,16,29). To identify additional nuclear-encoded components involved in mitochondrial double-strand break (DSB) repair, we screened EMS-mutagenized *Arabidopsis* plants for this phenotype to identify additional (nuclear-encoded) components of the DSB repair machinery. One of the mutants identified carried a missense mutation in SUV3 (Suppressor of Var1 3), a gene encoding a mitochondrial ATP-dependent 3′→5′ RNA helicase. The yeast and human homologs of SUV3 function as a core component of the degradosome, associating with mitochondrial RNA exonucleases and unwinding RNA substrates to facilitate their degradation (30–32). In contrast, the function of SUV3 in plants remains poorly understood. Early studies demonstrated that *Arabidopsis* SUV3 (AtSUV3) localizes to mitochondria and exhibits ATPase activity (33). In rice, SUV3 (OsSUV3) is associated with salinity stress tolerance and displays bidirectional helicase activity (33–35). By examining the effects of ciprofloxacin-induced mitochondrial DNA damage, here we demonstrate that *Arabidopsis suv3^-^*mutants are defective in mitochondrial DNA repair. Furthermore, we show that AtSUV3 is an ATP-dependent RNA helicase with 3′→5′ directionality and propose a mechanism by which SUV3 may contribute to the repair of mitochondrial double-strand breaks (DSBs). Collectively, our findings identify AtSUV3 as a previously unrecognized factor involved in mitochondrial DSB repair and expand our understanding of how mitochondrial genome maintenance mechanisms have diversified in plants.

## MATERIALS AND METHODS

### Ethylmethanesulfonate mutagenesis

Approximately 5,000 *Arabidopsis thaliana* seeds (100 mg) were mutagenized with ethylmethanesulfonate (EMS) following previously described protocols (36,37). Seeds were treated in EMS solutions of 0.15%, 0.20%, or 0.25% for 15 h, rinsed thoroughly, and sown on soil. For each of the three EMS concentrations, ∼1,000 mutagenized seeds were sown on four trays of soil. Trays were stratified for 2 days at 4 °C and transferred to a growth chamber maintained at 22 °C under a 16-h light/8-h dark photoperiod. M2 progeny seeds from each tray were harvested, pooled, and assigned pool designations: A–D for 0.15% EMS, E–H for 0.20% EMS, and J–M for 0.25% EMS. From each pool, 200–400 M2 individuals were grown under the same conditions, self-pollinated, and their seeds collected. Resulting M3 seed families were labeled according to their M2 pool of origin and a sequential number plant identifier.

### Screening candidates

Collected M3 seeds were surface sterilized using chlorine gas (Lindsey, Rivero et al. 2017) and plated on 1× Murashige and Skoog medium supplemented with Gamborg’s vitamins (Sigma), 10 g/L sucrose, 5 mg/L nystatin dehydrate (Indofine), and 8 g/L plant tissue culture agar (Carolina Biological Supply). The medium was adjusted to pH 5.7 (hereafter called MS plates). To screen for ciprofloxacin (Cipro) hypersensitivity, plates were supplemented with 0.5 µM ciprofloxacin (CIP plates), prepared from a 5 mM stock solution dissolved in 5 mg/mL MES hydrate, pH 5.7.

Approximately 12 sterilized M3 seeds from each family and 3 Col-0 seeds as wild-type controls were plated per CIP plate. Seeds were stratified at 4 °C for 2 days, then transferred to a growth chamber set at 25 °C with a 12-h light/12-h dark photoperiod for germination and growth. Each putative mutant was categorized according to its growth phenotype, with confirmatory rescreening on CIP plates when required, and parallel testing on MS plates to validate phenotypes initially observed under cipro exposure. Depending on the growth level on CIP plates, mutants were subdivided into two categories: all-dead and mostly-dead (38). The all-dead plants failed to develop beyond germination. In contrast, mostly-dead plants displayed enough early growth to reach the 2-4 leaf stage, and when transferred to MS plates, frequently recovered and resumed normal development. One M3 line called K147 was categorized as mostly-dead on ciprofloxacin and further characterized.

### Backcrossing and Illumina sequencing of EMS line

Re-screening M4 progeny from K147 on CIP plates confirmed homozygosity. Homozygous K147 plants were backcrossed to the parental Col-0 stock used for EMS mutagenesis. BC_1_F_1_ (first filial generation from a backcross) progeny was grown on soil and allowed to self-pollinate, producing BC_1_F_2_ seeds. BC_1_F_2_ seeds were germinated on CIP plates, and surviving seedlings were transferred to MS plates without CIP. DNA was extracted from seedlings that recovered, and Illumina sequencing libraries were prepared and sequenced as described in Rowan et al.(7)

### Sequence analysis of EMS line

Raw Illumina reads were aligned to the *Arabidopsis thaliana* TAIR10 reference genome using BWA-MEM (39) Variants were called with SAMtools mpileup and BCFtools (40) VCF files were converted to SHOREmap format and analyzed with SHOREmap annotate using TAIR10 GFF annotations (41,42). Alleles were filtered for EMS-consistent changes (G/C→A/T), location within coding regions, predicted nonsynonymous or nonsense effects, and allele frequency >0.5. Variants meeting these criteria were further validated against Col-0 genomic DNA and the backcrossed M3 mutant.

### T-DNA mutant line identification and selection

To complement the EMS forward-genetic approach and obtain an independent allelic perturbation of *AtSUV3*, an *AtSUV3* T-DNA insertion line (SALK_078210) was obtained from the Salk Institute Genomic Analysis Laboratory (43), and was used as a reverse-genetic tool to reduce *SUV3* function and test whether partial loss of *AtSUV3* recapitulates the ciprofloxacin hypersensitivity and genome instability phenotypes observed in the EMS mutant. This line contains a T-DNA insertion in exon 6 of the *AtSUV3* (At4g14790) coding sequence on chromosome 4. The insertion confers kanamycin resistance (Kan^R^) as a selectable marker. To identify plants carrying the T-DNA insertion, generation two (T₂) seeds were sown on 0.5× Murashige and Skoog (MS) medium supplemented with MES (Research Products International, SKU M70800-50.0) and 60 µg/mL kanamycin. After 7–10 days, surviving seedlings were transplanted to soil for growth under standard conditions.

### DNA extraction and genotyping of T-DNA mutant line

Genomic DNA was extracted from individual T₂ leaves using a hybrid cetyltrimethylammonium bromide (CTAB) and solid-phase reversible immobilization (SPRI) bead purification methods (44,45). PCR genotyping was performed using primers flanking the T-DNA insertion site and a right-border T-DNA primer. Amplification products of approximately 1,000 bp indicated the wild-type allele, while 470–780 bp products confirmed the presence of the T-DNA insertion. Heterozygous plants yielded both amplicons.

### gRNA design and cloning

To ensure efficient gene knockout, two gRNAs were designed to target *SUV3*, one in exon 3 and the other in exon 7. The gRNAs were generated using Geneious’ CRISPR Guide RNA Design Tool Software (https://www.geneious.com/crispr/). gRNAs were ordered in the pUC57 vector from GenScript along with the appropriate gRNA scaffolding and flanked by *Bbs*I sites. To transfer the gRNA construct into the entry vector, pENL1L2_Chimera, Golden Gate assembly was performed, and the resulting entry vector with gRNA was transformed into Top10 OneShot *E. coli* (ThermoFisher Scientific cat. no. C404010, Fauser et al. 2014, Bari et al. 2019). Successfully transformed *E. coli* colonies were used to extract the entry vector which was subsequently used to Gateway clone the gRNA construct into the pMR333 destination vector (provided by Dr. Mily Ron, University of California, Davis, CA, USA, Conklin et al. 2020). *E. coli* was transformed using the destination vector, confirmed with Eurofins Genomics Whole Plasmid Sequencing service, and the confirmed destination vector was used to transform *Agrobacterium tumefaciens* c58c1. Transformed *Agrobacterium* colonies were used to floral dip *A. thaliana* (46).

### Basta selection

Seeds from floral-dipped plants were collected and planted for Basta (Glufosinate Ammonium) treatment. Treatment included spraying 300 μM Basta (plantmedia.com, SKU# 30632007-5) directly onto the soil at the time of planting, at the time of first emergence, and one week after emergence. If the wild-type control survived the treatment after the third spray, more treatments were done until the wild-type control was affected. After treatment, surviving plants were allowed to grow until maturity, and seeds were collected.

### Microscopy

Seeds from plants surviving Basta treatment were screened for green-fluorescent protein (GFP) fluorescence using a Nikon SMZ25 stereo fluorescence microscope, as the destination vector includes a seed coat EGFP marker. Seeds that were positive for GFP were planted on soil and had leaves collected for DNA extraction. Extracted DNA was used to verify a deletion with PCR around the expected target sites and to confirm the presence of the CRISPR/Cas9 construct.

### T-DNA Insertion location analysis of CRISPR mutant (FPNI-PCR)

To determine the location of T-DNA sequence insertion in the nuclear genome, FPNI-PCR was performed following previous methods (Wang, Ye et al. 2011) with the primers included in **Table S2.**

### Whole genome sequencing and bioinformatics

Genomic DNA was extracted from T_2_ *suv3*^-^ CRISPR mutant and wild type plants using SPRI-bead based methods (Rowan, Seymour et al. 2017) and sent to SeqCenter (seqcenter.com, Pittsburgh PA, U.S.A.) for Illumina Whole Genome Sequencing. FASTA paired-end reads were interleaved and aligned using BWA-MEM (47). against a reference genome modified to exclude the mitochondrial genome copy on nuclear chromosome 2. Duplicate reads were removed, and the resulting BAM file was sorted and indexed with SAMtools. Samtools depth was used to make a text file containing the depth at each nucleotide for each chromosome. These depth values were then averaged to give the average depth on each nuclear chromosome, as well as the mitochondrial and chloroplast genomes.

Mitochondrial-enriched DNA was extracted from plants using previously described methods (48,49). DNA samples were sent to SeqCenter for Oxford Nanopore sequencing. Reads shorter than 1000bp were filtered out. Reference-based alignment was performed with minimap2. Reads that mapped to the mitochondrial genome with minimap2 were extracted and used for *de novo* assembly of the mitochondrial genome using Flye, and the resulting genome structure was visualized with Bandage (50,51). Structural variants were called using the SVIM software (52).

### Germination rate

10 seeds of wild type, 10 T_2_ *suv3^-^* mutant seeds, and 10 T_3_ *suv3*^-^ mutant seeds were sterilized and plated on 0.5× MSA. This was replicated 4 times. After vernalizing for 2 days at 4°C, the plates were moved to a growth chamber with 16 hours light, 8 hours dark at 25 °C. The number of seeds that germinated was counted 4 days post-vernalization.

### Root length assay

All seeds were surface sterilized with chlorine gas as described above. Seeds were plated on 9x9 cm square plates containing 0.5× MSA (pH 5.7) both with and without the addition of 0.5 μM ciprofloxacin. After 2 days of vernalization, they were moved to the growth chamber where they were kept at a roughly 70° angle. Plates were imaged 9 days later using the brightfield setting on a GelDoc imager. Root lengths were measured with the flexible line tool in ImageJ software (53).

### Silique length analysis

Green siliques from 41-day old plants were imaged under a dissecting microscope. Thirty siliques of each genotype were measured, from 3 different plants (10 siliques each), using ImageJ software (53). The difference in silique length was statistically analyzed using a 2-sample t-test.

### Salt sensitivity assay

Mutant and wild-type seeds were surface-sterilized using chlorine gas (described previously) and plated on 0.5× MSA (0.6% agar, pH 5.6, 0.86% sucrose) plates supplemented with 0 mM, 150 mM, 200 mM, or 250 mM NaCl. The plates were vernalized for 2 days at 4℃ and then moved to the growth chamber. The number of seeds that germinated was counted 22 days after sowing.

### Heat sensitivity assay

Mutant and wild type seeds were surface sterilized using chlorine gas and plated on 0.5× MSA plates (1% agar, pH 5.6) with 15 seeds plated per genotype on each of 3 replicate plates. The plates were vernalized for 2 days at 4 °C and then moved to the growth chamber. Twelve days after sowing, 2 of the plates were subjected to a 3-hour heat treatment at 37°C in the dark. To control for the lack of light during the 3-hour incubation, the control plate was kept at room temperature in the dark for the same duration. After treatment, the plates were moved back to the growth chamber. The seedlings were monitored in the following days for bleaching and seedling death.

### Quantitative PCR analysis of mitochondrial repeat copy number

To measure repeat recombination in induced *suv3^-^* plant DNA, 2 sets of primers were designed that flank repeat L, copy #1 and repeat L, copy #2 (L1 and L2, respectively). The mitochondrial gene, *rrn18* was used as a reference gene for normalization, and wild type plants were used as a control. Primers are listed in **Table S2**. Reactions were set up in clear-bottomed 96-well plates. Each reaction was performed in both technical and biological triplicates. Each well contained 7μL SSOAdvanced Universal SYBR Green Supermix (Bio-Rad), 3 μL of each primer (2.5 μM), and 2 μL of DNA template normalized to a concentration of 0.5 ng/μL as measured by Qubit. Reactions were performed in CFX96 Real-Time System thermal cycler (Bio-Rad). Following an initial 5-minute denaturation step PCR was done for 45 cycles of 95°C for 5 min, 61°C for 30 sec, 72°C for 17 sec, plate read, followed by a melt curve from 65°C to 95°C in 0.5°C increments with a plate read.

Results were analyzed using the double delta method (54). Outliers from triplicates causing the standard deviation to be over 1.0 were removed and the average Cq was recalculated. The Cq value for each sample reaction was normalized to *cox1* by subtracting the *rrn18* Cq value from the Cq value of the same sample (ΔCq). Then, the Cq values were normalized to the wild type Cq values by subtracting the wild type Cq value from the induced samples from the same primer amplification ( ΔΔCq). ‘Relative expression’ of each repeat was calculated by the equation: 2^-( ΔΔCq). To check for significant differences in the ‘relative expression’ of each repeat form as compared to wild type, an unpaired t-test on the Δ Cq values was done.

### Expression and purification of AtSUV3

The CDS of *AtSUV3* (residues 40-1725) was cloned into pET-SUMO with restriction enzymes *Nde* I and *Bam* HI. Recombinant *AtSUV3* was expressed using the *Escherichia coli* BL21 C41 strain. Cell cultures were grown until they reached an OD_600_ of 0.8 and were induced with 1 mM IPTG (isopropyl-1-thio-β-D-galactopyranoside) at 16 °Cfor 18 h. Cells were recollected by centrifugation at 4 °C and resuspended in 30 mL lysis buffer (25 mM Tris-HCl pH 8.8, 1 M NaCl, 10 mM imidazole, 10% glycerol, 1 mM PMSF and 0.1% Triton X-100). The lysate was sonicated for 10 min at power 5 in a 5 s ON/5 s OFF cycle on ice in a Misonix Ultrasonic Procesor XL and centrifuged at 15000 rpm/40 min. The clear lysate was treated with PEI 0.02% (w/v) for 30 min and centrifuged at 15000 rpm/30 min, filtrated and loaded onto a Ni-NTA column. The column was washed with lysis buffer containing 30 mM imidazole. Proteins were eluted using lysis buffer supplemented with 500 mM imidazole and dialyzed in buffer (25 mM Tris-HCl pH 8.8, 300 mM NaCl, 2 mM EDTA, 10% glycerol and 5 mM DTT). *AtSUV3* was loaded into a Hi-Trap heparin column and further purified using a 0.1 M to 1 M NaCl gradient. Pure fractions were pooled and loaded into a Hi-Trap Q HP column and further purified using a 0.1 M to 1 M NaCl gradient. Pure fractions were pooled and concentrated then applied to a Superose™ 12 10/300 GL column in an ÄKTA pure™ chromatography system. Sample was run at 0.25 mL/min using 25 mM Tris-HCl 8.8, 1 M NaCl, 2 mM EDTA and 5 mM DTT as mobile phase. Elution was monitored by measuring absorbance at 280 nm.

### Oligonucleotide substrates

The oligonucleotide sequences used for cloning and biochemical assays are shown in **Table S1**. Oligonucleotides were purchased from IDT™. Substrates used for the helicase and binding assays were prepared by mixing 5’-end Cy5-labeled 18nt DNA with an equimolar amount of 34nt RNA or DNA in annealing buffer (10 mM Tris–HCl pH 8.0, 100 mM NaCl, and 1 mM EDTA) and then heated to 95 °C for 5 min followed by slow cooling at room temperature. Similarly, to generate transcription fork and splay duplex substrates, 5’-end FAM-labeled 50nt DNA was annealed with an equimolar amount of 33nt RNA and 50nt DNA. Also, dsDNA substrate, 3’-end Cy5-labeled 50-nts DNA was prepared using the same procedure with an equimolar amount of 22-nts DNA.

### Helicase activity assays

Unwinding assays were performed at 37 °C in reaction buffer (20 mM HEPES pH 7.5, 10 mM KCl, 1 mM DTT, 0.2 mg/ml BSA and 2.5 nM annealed substrates). Reactions were incubated with increasing concentrations of *AtSUV3* for 5 min at 4 °C, and the unwinding reactions were initiated by adding 10 mM MgCl_2_ and 5 mM ATP at 37 °C. Reactions were quenched after 30 min by adding 10 µL stop buffer 2x (8% Glycerol, 0.6% SDS, 40 mM EDTA). All samples were resolved in 12% native-PAGE TBE gel. Gels were visualized in an Amersham™ Typhoon™ biomolecular imager in the Cy5 or Cy2 channels.

### Nucleic acid substrate binding assays

To investigate *AtSUV3* binding, we used the substrates mentioned above. Reactions were incubated in reaction buffer (20 mM Tris-HCl pH 7.5, 20 mM KCl, 1 mM DTT, 10% glycerol, 2 mM EDTA, 0.1 mg/ml BSA and 1 nM annealed substrate) with increasing concentrations of *AtSUV3* for 30 min at 23°C. All samples were resolved in 12% native-PAGE TBE gel. Gels were visualized in an Amersham™ Typhoon™ biomolecular imager in the Cy5 channel.

## RESULTS

### Identification of *SUV3* mutants in Arabidopsis

From a pool of EMS-mutagenized *Arabidopsis thaliana* Col-0 plants, we isolated a mutant exhibiting a semi-dominant, variably expressive lethal phenotype on ciprofloxacin-containing media (**Fig. 1A**). The phenotype of this line, designated K147, was complicated as the line exhibited reduced seed viability on 0.5× MS even in the absence of ciprofloxacin, together with mild ciprofloxacin sensitivity in the parental Col-0 background. The K147 line was backcrossed to Col-0 and the F_1_ was self-crossed for segregant analysis by sequencing. Seeds were planted on agar media containing 0.5 mM ciprofloxacin. Approximately one quarter of the seedlings had either no true leaves or dying true leaves after 1 week. Those seedlings were transplanted to 0.5× MS media without ciprofloxacin and allowed to recover. Of those, typically 60-70% would recover. Those seedlings that grew to the 6 to 8 leaf stage were homogenized, and DNA was extracted and sent for Illumina sequencing. Briefly, DNA sequencing revealed a spike in allele frequency corresponding to a GC-to-AT transition in the gene annotated as At4g14790. This gene was subsequently identified as the *Arabidopsis thaliana* homolog of SUV3, a gene that has been extensively studied in *Homo sapiens*, and yeast as the helicase component of the mitochondrial RNA degradosome and in rice as a bidirectional helicase involved in the response to abiotic stress (**Fig.1B**). (35,55–57). Although the At4g14790 gene in *Arabidopsis* has been identified as a putative mitochondrial DExH-box RNA helicase with ATPase activity and with high homology to SUV3 proteins (56), its physiological role and *in vitro* activity has remained unknown.

**Fig. 1.**
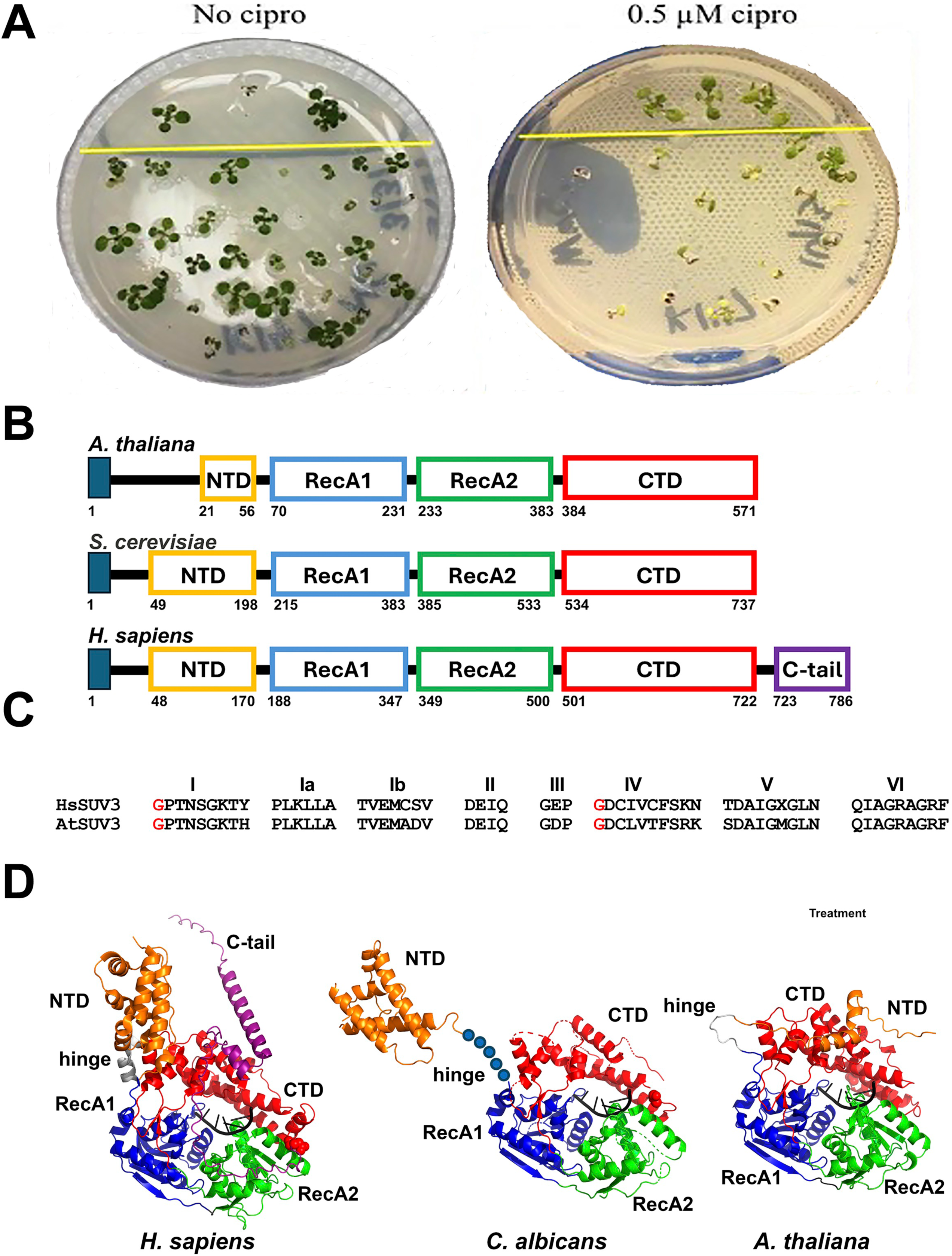
The ciprofloxacin-sensitive phenotype of the EMS mutant K147 results from disruption of the conserved mitochondrial helicase AtSUV3. **A)** Ciprofloxacin-sensitive phenotype of the EMS mutant K147. Representative M3 seedlings of the EMS mutant K147 grown on MS medium with (left) or without (right) 0.5 µM ciprofloxacin (CIP). On CIP-containing medium, K147 seedlings (left-below the yellow line) exhibit growth inhibition and chlorosis relative to the wild-type Col-0 controls (left-above the yellow line), consistent with impaired mitochondrial genome maintenance. This sensitivity motivated subsequent backcrossing and genetic analysis of the mutation. **C)** Structural comparison of SUV3 helicases from yeast, humans, and Arabidopsis and conserved RecA-like helicase motifs characteristic of Superfamily 2 helicases. Domain organization was annotated based on the crystal structure of human SUV3 as follows: N-terminal domain (NTD), gold; RecA1, blue; RecA2, green; C-terminal domain (CTD), red; and the human-specific C-terminal extension, purple. Structurally aligned sequences spanning the conserved helicase motifs of SUV3 proteins from diverse organisms are also shown. Residues predicted to interact with ATP are highlighted in red, residues involved in nucleic acid binding are shown in blue, and residues that stack with the adenine base are indicated in green (59). **D)** Structural model of AtSUV3 and comparison with yeast (*Candida albicans*) and human homologs. The human and yeast structures correspond to PDB entries 3RC8 and 6F4A, respectively. The human C-terminal tail (residues 716–786) is represented using an AlphaFold model.

*Arabidopsis* Suv3, as its human and yeast orthologs, contain a conserved SF2 helicase core that consist of two-tandem RecA1-like and RecA2-like domains and N-terminal and C-terminal extensions (58) (**Fig.1B and Fig. 1S**). AtSUV3 harbors the conserved RecA-like motifs required for NTP-dependent RNA unwinding and remodeling. In addition, AtSUV3 possesses the eight distinctive ATP-and RNA-binding motifs, including a DEIQ sequence in motif II instead of the canonical DExH motif, that set SUV3 proteins apart from other members of the SF2 helicase superfamily (**Fig.1C)** (59). A notable difference between human, yeast, and plant SUV3 proteins is the length of their N-terminal domain (NTD), which comprises approximately 148 amino acids in yeast, but only 28 and 35 residues in *O. sativa* and *A. thaliana*, respectively (**Fig.1B**). Thus, the NTD of *Arabidopsis* SUV3 is predicted to consist of only two short α-helices rather than a structured domain that adopts multiple conformations as in yeast and human SUV3 proteins (**Fig. 1C**). In addition, human SUV3 harbors an extended C-terminal tail (CTT) possibly involved in dimerization, that is absent in yeast and plant SUV3 (**Fig. 1S and 1D**).

The phenotype of the EMS allele (referred herein as *suv3^K147^*) was caused by a G to A transition that introduces the amino acid change **G252R** in Arabidopsis SUV3. The phenotype of the *suv3K147* allele was complex because it is semidominant, variably expressive, and exhibited partial recessive lethality. The G252R substitution in Arabidopsis SUV3 lies within motif IV. Structural and biochemical studies of human and fungal SUV3 proteins shown that motif IV contributes to RNA binding and the corresponding Gly residue in *Candida* and human SUV3 are completely buried within the hydrophobic core of the enzyme (55,58,60). Thus, the substitution of Glycine 252 with Arginine (AtSUV3-G252R) is predicted to substantially destabilize AtSUV3 **(Fig. 2A and B)**. *In silico* stability analysis using the program DUET on the AlphaFold model of AtSUV3 predicts that the G252R substitution has a strongly destabilizing effect (61), that is consistent with the physicochemical differences between the two residues: glycine consists of a hydrogen atom side chain, in stark contrast to the large, positively charged guanidinium group of arginine. The replacement of glycine 252 with arginine introduces both steric bulk and a charged group at a position that may perturb, ordisrupt, the hydrophobic core of AtSUV3 **(Fig. 2A and B)**, albeit molecular dynamics simulation show repacking of the side chains that interact with Gly or Arg at position 252 **(Fig. 2A and B)**, these structural changes are predicted to compromise the stability and structural integrity of AtSUV3-G252R and, consequently, its function.To substantiate this prediction, we estimated the changes in protein stability (ΔΔG) using DynaMut2, mCSM, SDM, and DUET programs for the corresponding glycine residue, based on the crystal structures of SUV3 from Arabidopsis, Candida and human (61,62)(**Fig. 2C**). The ΔΔG values range from −0.82 to −3.37 kcal/mol across the different experimental structures. However, all of them indicate a clear destabilizing effect of the mutation on diverse SUV3 orthologs.

**Fig. 2.**
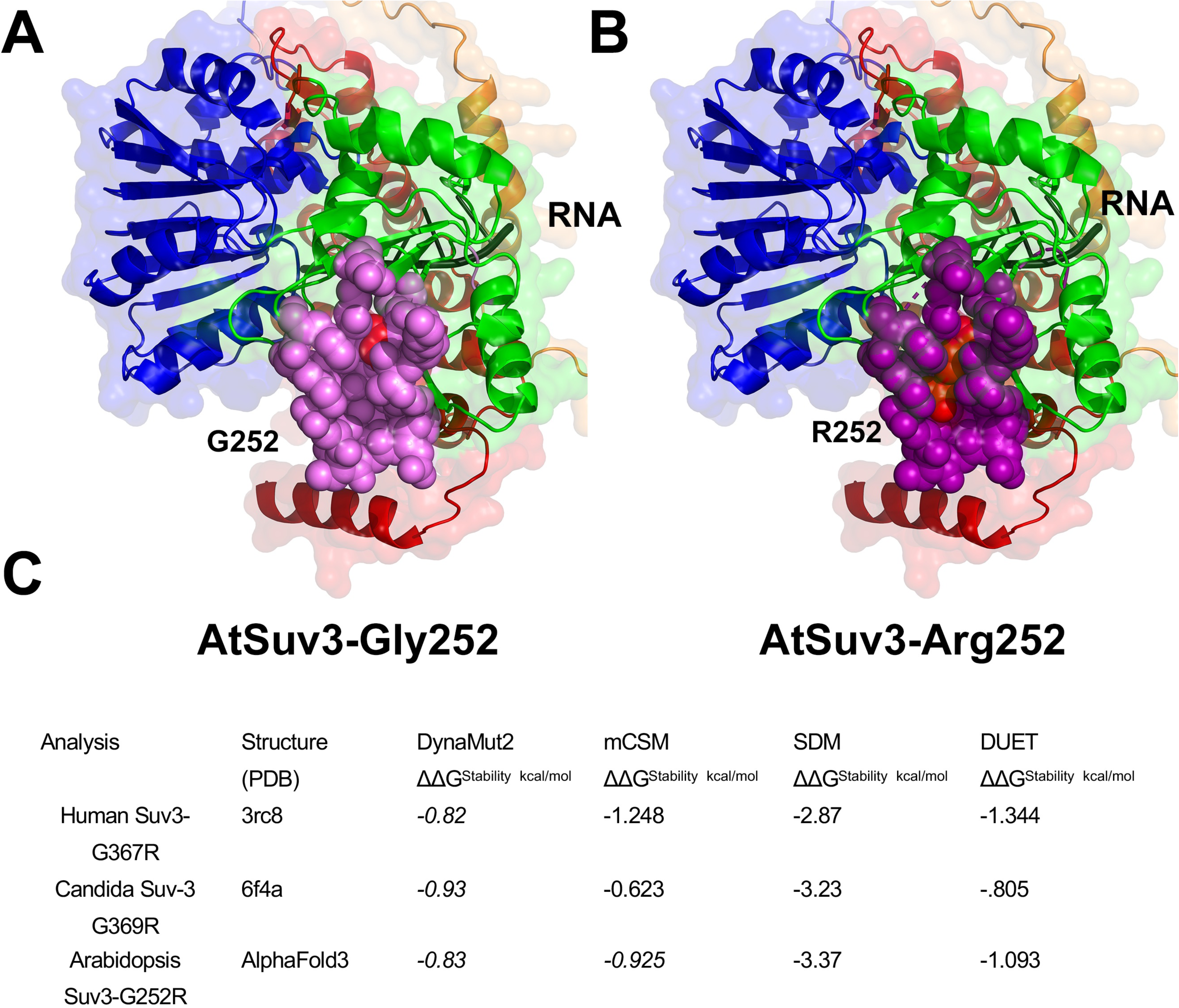
Structural consequences of the G252R substitution in AtSUV3. **A-B)** Structural model of AtSUV3 showing the location of Gly252 and its corresponding mtuation to Arg within the conserved helicase core. The structural model AtSUV3-G252R variant illustrating the replacement of glycine by a bulky, positively charged arginine side chain, which is predicted to alter local packing and intramolecular interactions. Residues of interest are shown as stick representations to emphasize the local structural environment surrounding the mutation site. Residues of interest are shown as spheres to highlight the local structural environment surrounding the mutation site. **C)** ΔΔG values for the mutation corresponding to G252R in *Arabidopsis*, human, and *Candida* SUV3, calculated with four different prediction methods.

### Loss of AtSUV3 does not affect plant growth, germination, root elongation, or reproductive development under standard growth conditions

To better characterize *SUV3* genetically, we obtained a T-DNA insertion mutation (SALK_078210) from the Salk Institute Genomic Analysis Library (43) and simultaneously constructed a CRISPR/Cas9 deletion mutation in *SUV3*. The T-DNA insertion in line SALK_078210 is located in exon 6 of the AtSUV3 (At4g14790) coding sequence on chromosome 4. However, this line was difficult to reliably genotype by PCR, possibly because of additional T-DNA fragments in the genome, and we were unable to isolate homozygous plants (**Fig. 3A**). For this reason, the CRISPR-induced deletion was solely used for further studies. Plants successfully transformed with the Cas9 and gRNA construct produced progeny with seed coat GFP in a roughly 50/50 ratio, suggesting a hemizygous insertion of the CRISPR construct in the nucleus. A *suv3^-^*deletion was verified using the results of Illumina sequencing. The gRNA target in exon 3 caused a large, 417bp deletion, which removed the gRNA site, as well as the entirety of the exon. Notably, this deletion removed the predicted ATP binding site of the protein, as the deletion was a frameshift followed by an insertion of eight incorrect amino acids followed by a stop codon. The gRNA site in exon 7 had a single base pair deletion, downstream of multiple in-frame stop codons that is predicted to given rise to a nonfunctional SUV3 protein. To confirm that any effect of a CRISPR deletion mutant is indeed due to the truncation of the gene of interest, and not due to a disruption of a gene with the nuclear T-DNA insertion, we determined exactly where the T-DNA insertion occurred within the nuclear genome, using FPNI-PCR. The insertion site of the T-DNA sequence in the nuclear chromosome was found to be on chromosome 5 in an area that does not disrupt any annotated genes or features in the reference genome (nuclear chromosome 5, base 26,137,967). This result was confirmed in the Illumina assembly, with supplemental reads at that location mapping to T-DNA sequence.

**Figure 3:**
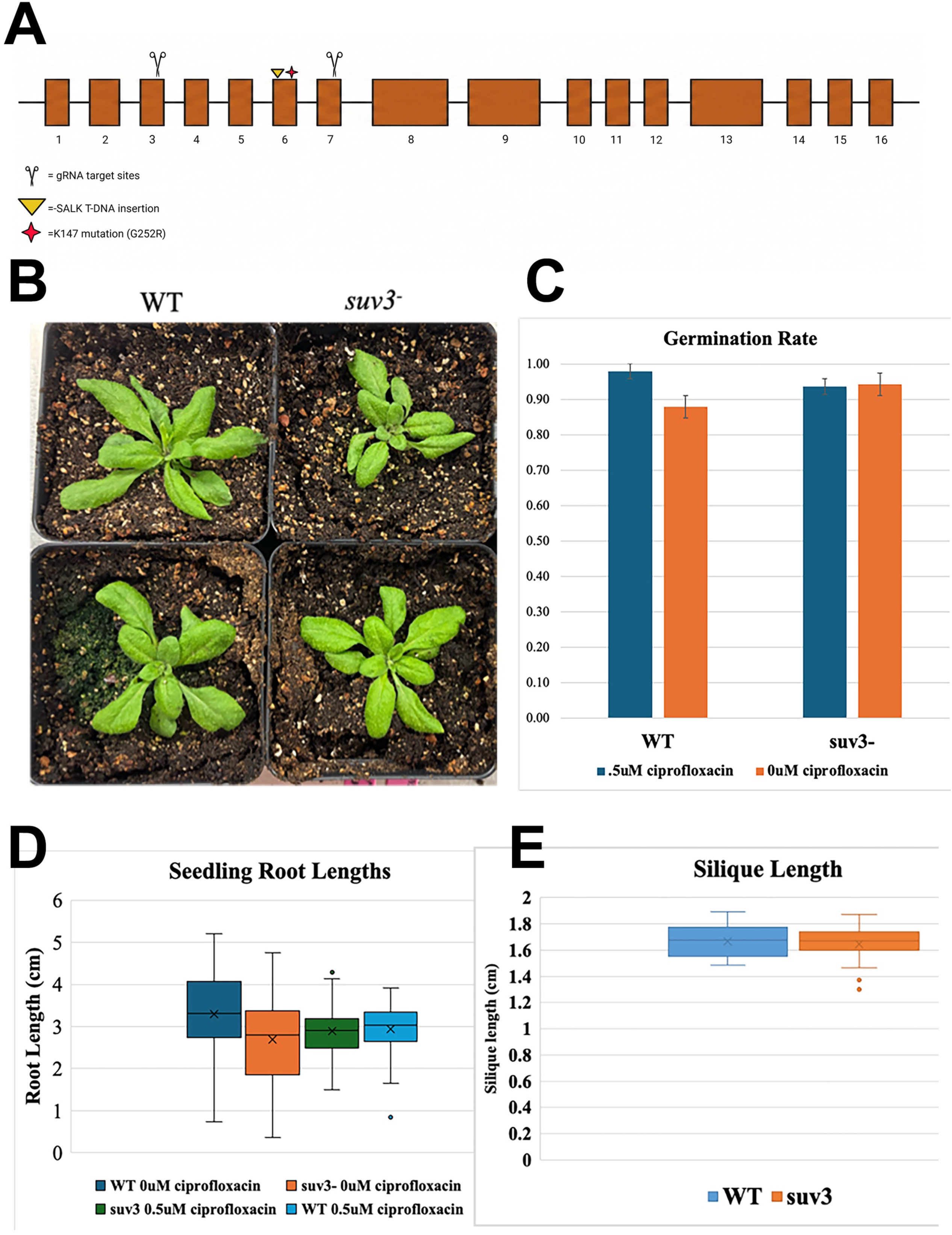
The *suv3^-^*mutant exhibits no detectable developmental defects or increased sensitivity to ciprofloxacin. **A) Fig. 2 Generation of the *suv3* knockout mutant.** (a) Schematic of the *AtSUV3* locus showing the position of the SALK_078210 T-DNA insertion and the CRISPR/Cas9 target sites. The CRISPR-generated *suv3* allele contains a 417-bp deletion spanning exon 3, resulting in loss of the predicted ATP-binding region, a frameshift, and a premature stop codon; a 1-bp deletion at the exon 7 target site is also indicated. **B)** Growth of *suv3^-^* mutant seedlings on soil compared to wild type (Col-0). Photo taken 21 days after initial planting. **C)**. Germination rate of wild type seeds compared to *suv3^-^*seeds on 0.5x MSA plates with or without ciprofloxacin. *suv3^-^*n=120 each treatment, WT n=40 each treatment. No significant difference between either comparison (WT vs *suv3^-^* ciprofloxacin treatment or *suv3^-^*with ciprofloxacin vs *suv3^-^* without ciprofloxacin, as measured by a t-test). **D)** Root lengths of 9-day old seedlings both with and without ciprofloxacin (0.5 μM) treatment. No significant difference between wild type (WT) and *suv3^-^*mutant seedlings both with and without treatment. (WT 0 μM ciprofloxacin vs *suv3^-^* ciprofloxacin p-value=0.023, WT 0.5 μM ciprofloxacin vs *suv3^-^* 0.5 μM ciprofloxacin p-value=0.604, as measured by t-test). WT 0 μM ciprofloxacin n=43, *suv3^-^*-0 μM ciprofloxacin n=29, *suv3^-^*0.5 μM ciprofloxacin n=84, WT 0.5 μM ciprofloxacin n=89. **E)** Mature green siliques from 41-day old plants were measured in both wild type and *suv3^-^* plants. No significant difference in silique lengths (p-value=0.54, t-Test: two-sample assuming equal variance, n=30 each genotype).

When allowed to grow without any stress or treatment, *suv3*^-^mutants grew normally as evidenced by their rosette size on soil and normal germination rate on 0.5× MS plates (**Fig. 3B, 3C**). Root growth measurements of seedlings, which is often used to approximate mitochondrial health (29,63), showed no difference in seedling root length between wild type and *suv3^-^*, both with and without ciprofloxacin treatment (64) (**Fig. 3D**). Siliques of 41-day-old plants were measured and showed no difference in length between wild type and *suv3^-^* (**Fig. 3E**). Overall, no difference was seen between the mutant and wild type plants under normal growth conditions.

### Mitochondrial genome structure in CRISPR-induced *suv3^-^* mutants

#### Mitochondrial DNA copy number

The average sequencing depth of each nuclear chromosome was compared with that of the mitochondrial genome to estimate relative mtDNA copy number. In the *suv3*^-^mutant, the relative mtDNA copy number was 9.97, whereas in four independently sequenced wild-type samples it ranged from 15.8 to 24.5 (**Table 1**). In *Saccharomyces cerevisiae*, suv3 mutants exhibit a severe reduction or complete loss of mitochondrial DNA (57). Thus, the decreased relative mtDNA copy number observed in the *Arabidopsis suv3*^-^ mutant suggests that SUV3 contributes to the maintenance of mitochondrial genome copy number in plants, consistent with its proposed role in yeast.

**Table 1.**
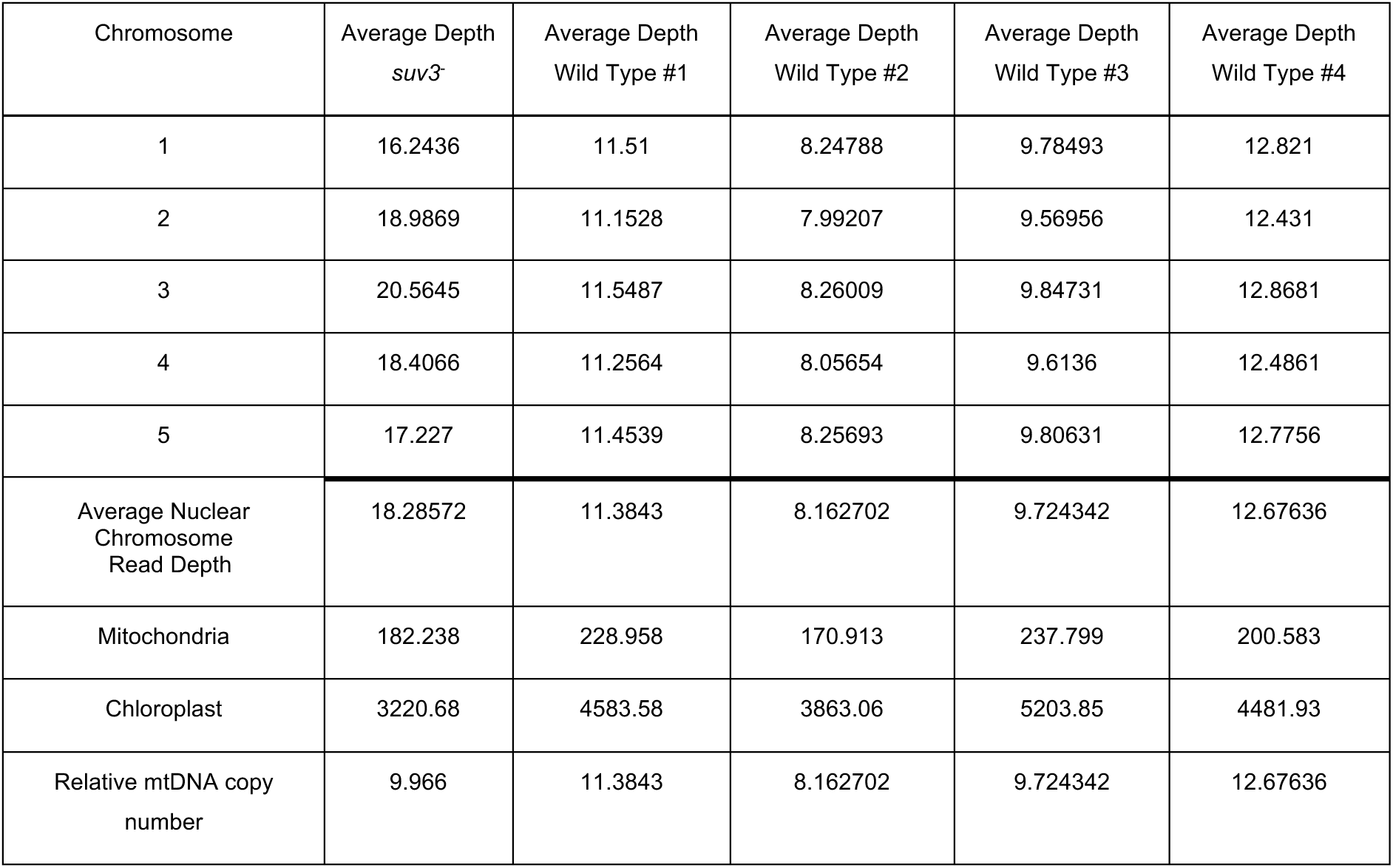
Read depth comparison and relative mtDNA copy number between suv3-mutant and wild-type Arabidopsis.

#### Mitochondrial genome organization

*De novo* assembly of mitochondrial reads of the *suv3^-^*mutant showed a mostly wild-type consensus structure, with the four main segments of the genome being separated by the large repeats **(Fig. 4A).** However, the reference-based assembly using the Nanopore long reads showed a large increase in structural variants compared to the Col-0 (wild type) Nanopore sequenced sample (**Fig. 4B**).

**Figure 4:**
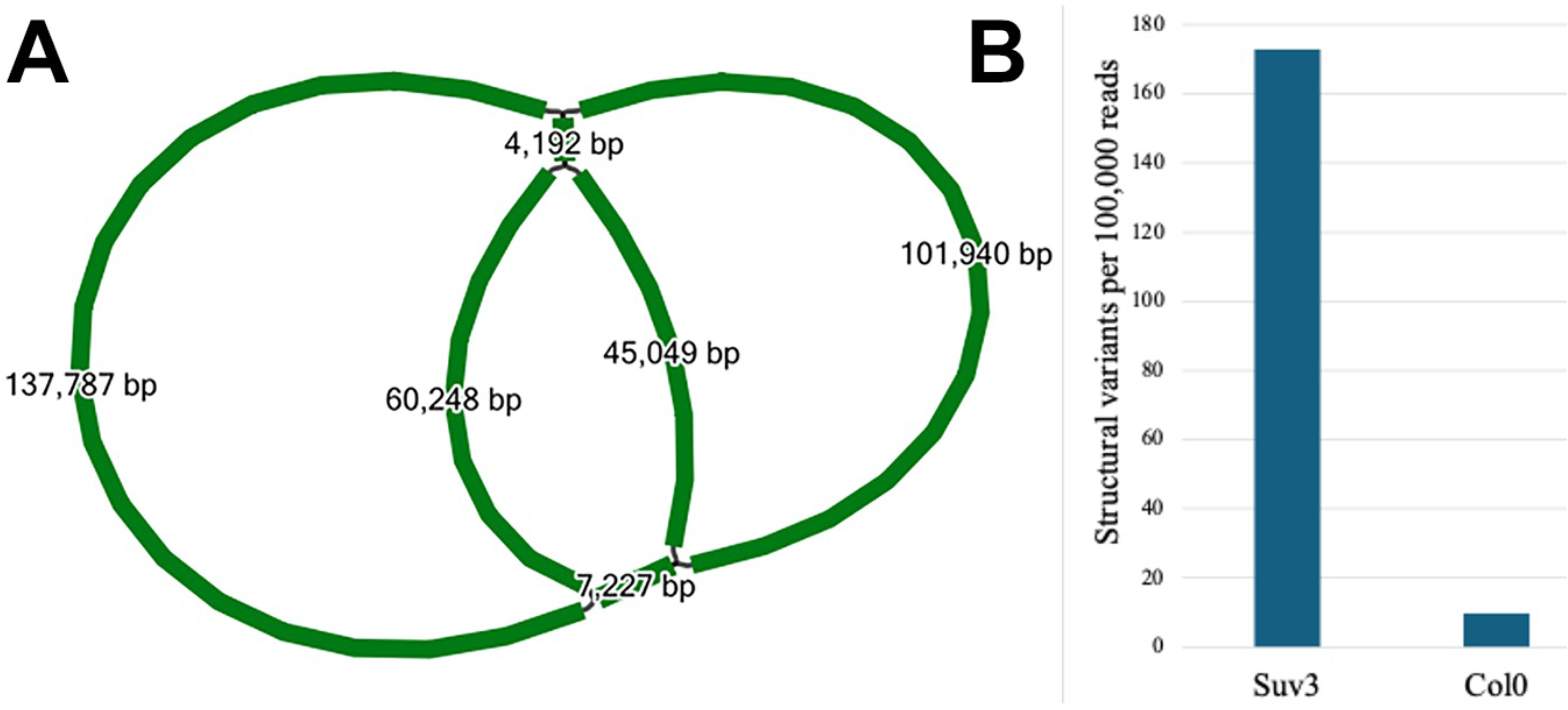
**A)** *De novo* assembly graph of *suv3^-^* mutant mitochondria. Assembly was performed using Flye and the resulting .gfa file viewed in Bandage. Fragments are labeled by length. Two small fragments (7,227bp and 4,192bp) are the Large #1 and Large #2 repeats, respectively. **B)** Structural variant counts in the mitochondrial genome of *suv3^-^* compared to wild type (Col-0). Structural variants called using SVIM from the minimap2 reference-based alignment and normalized to the number of structural variants per every 100,000 reads.

### AtSUV3 contributes to stress tolerance and maintenance of mitochondrial genome stability

The *suv3*^-^ mutants were shown to be highly salt sensitive, matching the results seen in rice *suv3*^-^ mutants (65) **(Fig. 5A).** The germination rate of the 22-day old *suv3*^-^ mutant seedlings was significantly lower than wild type on all concentrations of NaCl tested, with zero *suv3*^-^seeds germinating on the highest salt level. The *suv3*^-^ mutants were also shown to be much more sensitive to heat stress than wild type (**Fig.5B**). The mutant seeds bleached and died at a much faster rate than wild type (within 2 days post heat treatment). Both the wild type and mutant seeds on the untreated control plate had a 100% survival rate.

**Fig. 5:**
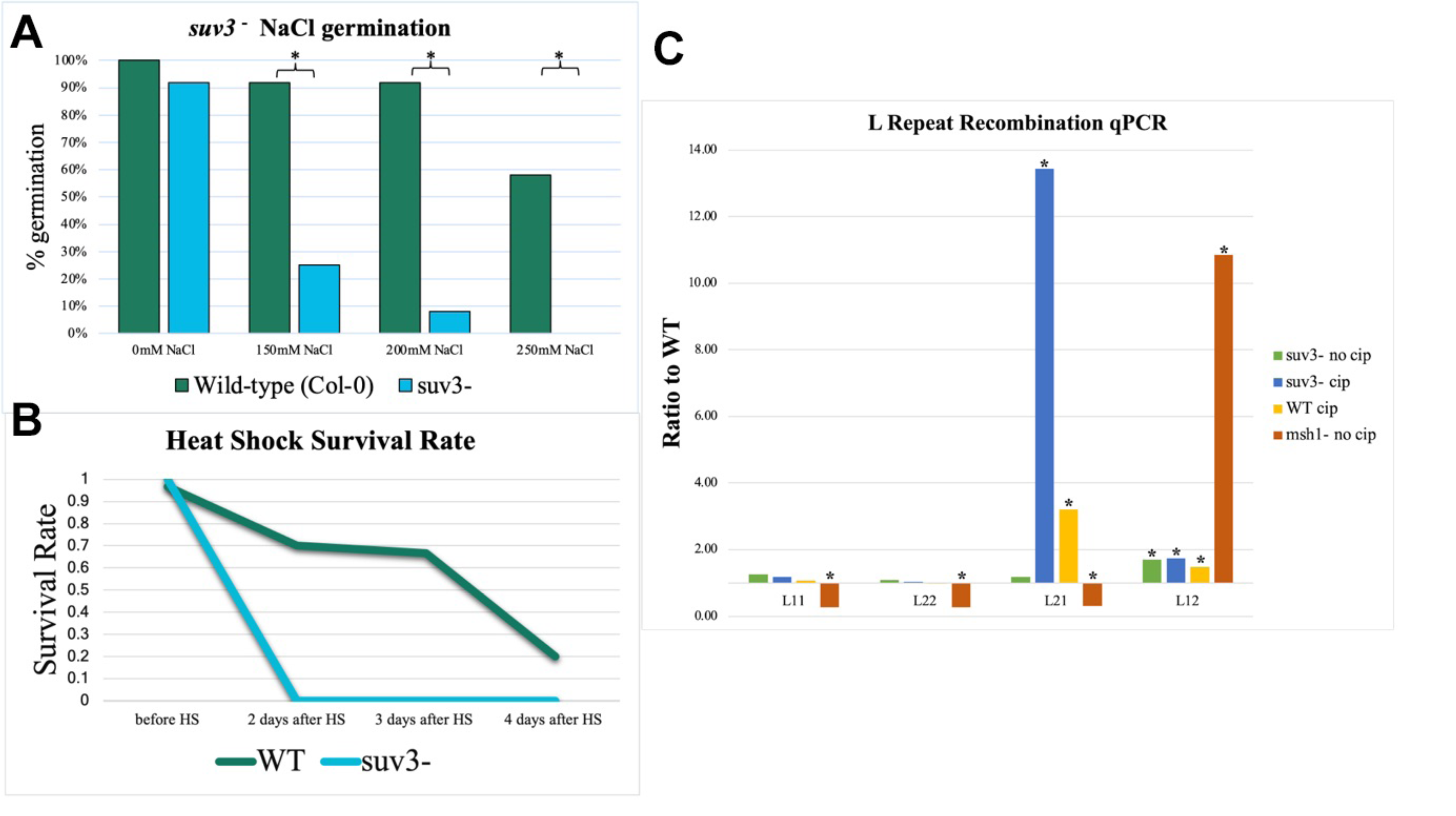
Stress responses and mitochondrial genome stability in the suv3 mutant. **A)** Percent germination of mutant and wild type (WT) seeds on plates containing varying levels of NaCl. n=12 seeds per genotype per treatment. (*=p< 0.05 Fisher’s Exact Test). **B)** Survival rate of wild type compared to *suv3*^-^ seedlings after a heat shock (37 °C, 3 hours) followed by recovery in the growth chamber. n=30 seedlings of each genotype. **C)** L repeat recombination qPCR on *suv3^-^* mutants with and without ciprofloxacin treatment (0.5 μM), compared to wild type on ciprofloxacin treatment. *msh1*^-^ results included as a point of comparison (three biological replicates, 1 technical replicate). All values normalized to *rrn18* and depicted as a ratio to wild type (no treatment) on the y-axis. *=p<0.05 (t-test on delta Cq values), n=3 biological replicates, 2 technical replicates.

### Repeat-mediated recombination

One common way of checking for mitochondrial genomic stress or instability is with qPCR analyses of intermediate-sized mitochondrial repeats (66). “Stress” in this context can be imposed by the effect of missing a DNA repair protein, such as Msh1, or by the application of genotoxic treatment such as ciprofloxacin (10,11,18,67). Whether it is caused by genotoxic treatment or mutations that compromise recombination and repair, stress leads to an increased frequency of recombination between intermediate sized repeat pairs, which can be quantified through qPCR. Recombinational repair can be accurate and leave no mutations through homologous recombination mechanisms, but it can also be very error prone, leaving expansions, and rearrangements due to the inaccurate mechanisms of recombination at ectopic sites. This can increase mitochondrial genome heteroplasmy, skewing towards the recombinant forms of the repeats. We measured repeat recombination at repeal L using qPCR in *suv3*^-^ mutants treated with and without ciprofloxacin. This repeat is typically used as a readout of disruption of genome maintenance processes in *Arabidopsis* mitochondria (6,68). CRISPR-induced *suv3*^-^ mutants without ciprofloxacin treatment show a small but consistent and significant increase in the recombinant form of the L repeat, L12. However, under ciprofloxacin treatment, the *suv3*^-^ mutants see a large and consistent increase in the recombinant form L21 (**Fig. 5C**). L repeat recombination is generally asymmetric, with the L12 recombinant form consistently enriched in DNA repair mutants (such as in *msh1*^-^) and after ciprofloxacin treatment. The increase in repeat form L21 is unique for the *suv3*^-^ mutant and suggests that although the Suv3 protein is playing a role in preventing ectopic recombination, it is doing so in a different way than other DNA repair proteins, such as Msh1.

### AtSUV3 unwinds dsDNA and dsDNA:RNA hybrids harboring 3′tails as a 3′ → 5′ helicase

Although several plant RNA and DNA helicases, including AtTwinkle, RadA, RecG, and the DExH-box RNA helicases ABA OVERLY SENSITIVE 6 (ABO6) and RH33, have been implicated in mitochondrial RNA splicing and genome maintenance relatively few have been biochemically characterized (13,29,69,70). To this goal we subcloned the SUV3 gene lacking its predicted mitochondrial targeting sequence (residues 1 to 21) as an N-terminal SUMO fusion protein. SUMO-tagged *AtSUV3* was purified through a three-step procedure consisting of immobilized metal affinity chromatography (IMAC), heparin chromatography, and size-exclusion chromatography (**Fig. 2SA**). Recombinant AtSUV3 (550 amino acids) has a predicted molecular mass of 61.3 kDa, which increases to approximately 74.7 kDa upon addition of the SUMO and His tags. After purification, a predominant protein band of 75 kDa was observed, consistent with the expected size of the fusion protein and initial helicase assays corroborated the capacity of *AtSUV3* for duplex unwinding with 3′→5′ polarity mainly under ATP-dependent conditions (56) (**Fig. 2SB and C**).

**Fig. 2S.**
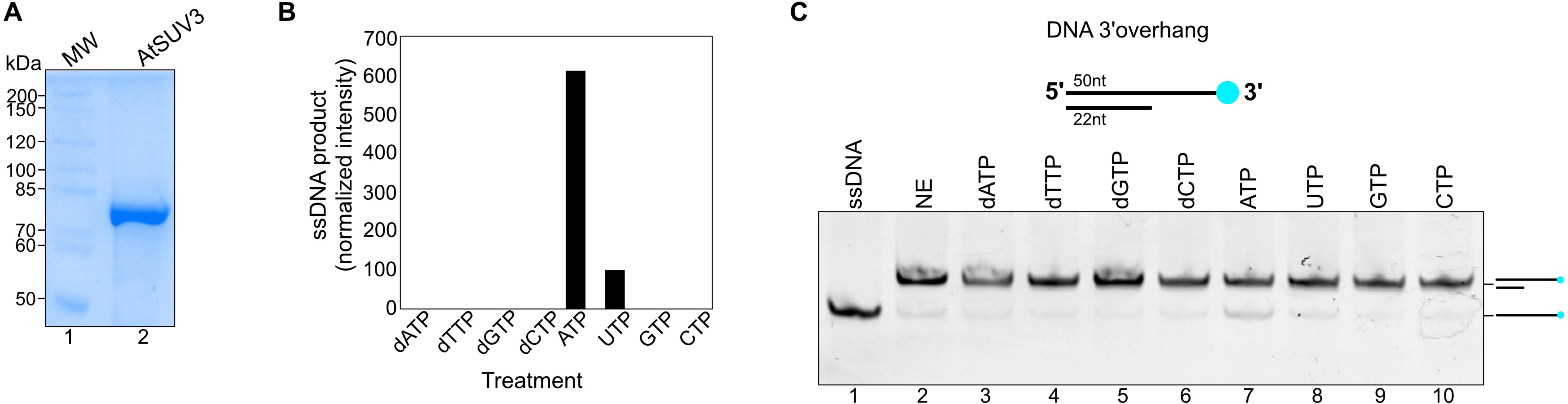
AtSUV3 is an ATP dependent 3′ → 5′ helicase. **A)** 10% SDS–PAGE showing the purified recombinantly expressed *AtSUV3* protein stained with Coomassie Blue. **B)** Helicase activity by *AtSUV3* requires ATP. Quantification of helicase activity represented as a bar graph depicts duplex unwinding under different nucleotides. Robust unwinding is observed only in the presence of ATP, whereas no detectable activity is measured in reactions lacking other nucleotides. **C)** Representative native gel showing the unwinding by AtSUV3 on a 3′overhang in comparison to a control with no enzyme (NE) (lane 2)

We therefore sought to biochemically characterize SUV3 from *Arabidopsis*. Human and yeast SUV3 exhibit strict unwinding with 3′→5′ directionality (71,72). This is in contrast to rice SUV3, which is reported to display dual unwinding polarity (35). Accordingly, we evaluated the binding and unwinding activities of *AtSUV3* using a panel of duplex substrates, including dsDNA, DNA–RNA hybrids, and DNA–DNA duplexes harboring either 5′ or 3′ single-stranded overhangs. Under our experimental conditions, *AtSUV3* exhibited selective binding exclusively to 3′ overhang substrates consisting of RNA:DNA hybrids with an RNA 3′ overhang (**Fig. 6 A-E**). Unwinding experiments showed an absolute requirement for a 3′ overhang indicating strict 3′→5′ polarity (**Fig. 6 F-J**). Notably, *AtSUV3* preferentially unwinds DNA–RNA hybrids in comparison to DNA–DNA duplexes, suggesting a preference for transcription-associated substrates (**Fig. 6I and 6J**). Thus, our biochemical result sharply contrasts with the dual substrate polarity established by rice SUV3 (OsSUV3) (35). This result is particularly noteworthy because AtSUV3 and OsSUV3 share more than 65% amino acid identity. Although the translocation polarity of human SUV3 was initially controversial, with early studies reporting 5′→3′ activity and subsequent work from the same group proposing an exclusive 3′→5′ directionality (71,73) the issue has since been definitively resolved. Multiple independent biochemical and structural studies have established that SUV3 helicases are bona fide 3′→5′ RNA helicases (58,74), indicating that this directionality is a conserved and fundamental property of the SUV3 family.

**Fig. 6:**
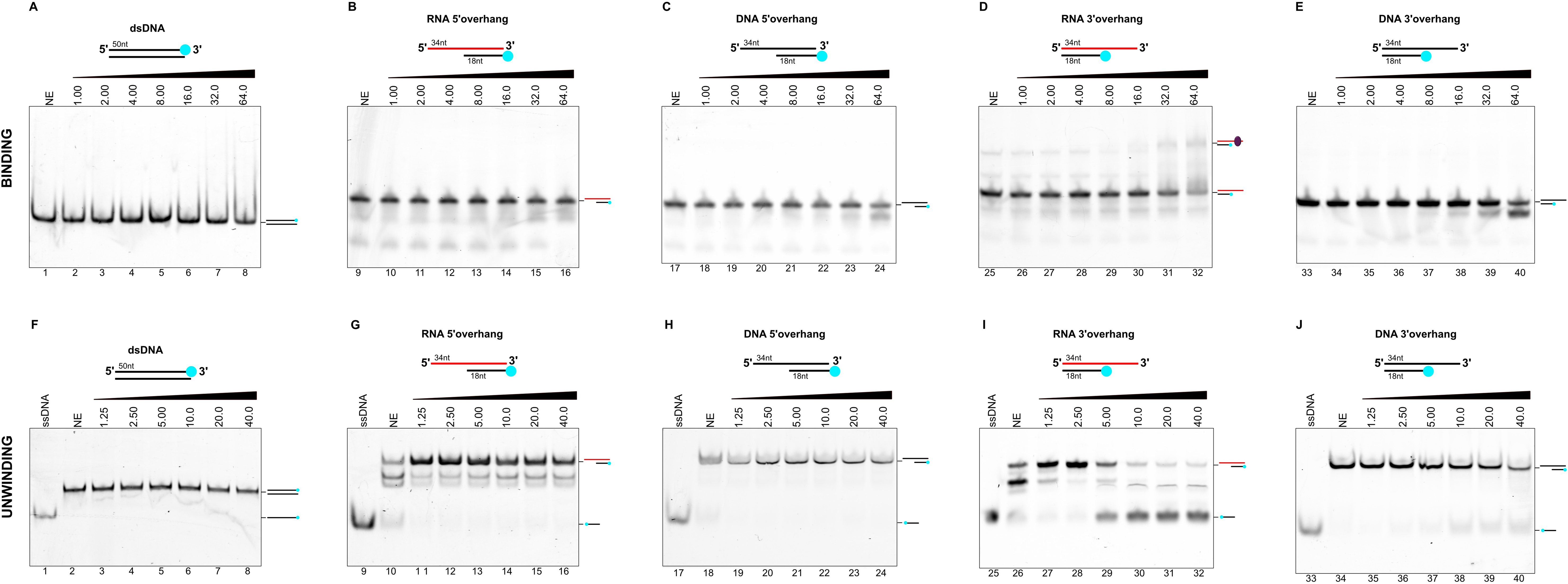
A*t*SUV3 is a DNA and RNA 3′ → 5′ helicase. **(A–E)** Electrophoretic mobility shift assays (EMSAs) assessing substrate binding by *AtSUV3* to substrates with distinct architectures: blunt-ended dsDNA **(A)**, RNA:DNA hybrid with a 5′ RNA overhang **(B)**, DNA duplex with a 5′ DNA overhang **(C)**, RNA–DNA hybrid with a 3′ RNA overhang **(D)**, and DNA duplex with a 3′ DNA overhang **(E)**. Increasing concentrations of *AtSUV3* (1.35–40 µM) were incubated with labeled substrates and resolved on native 10% polyacrylamide gels. **(F–J)** Helicase assays performed under ATP-dependent conditions using the same panel of substrates. Strand separation was monitored by the appearance of the labeled single-stranded substrate, which represents complete unwinding, whereas the no-enzyme (NE) control shows intact duplex substrate. No helicase activity is detected on blunt-ended substrates **(F)** or on duplexes containing 5′ overhangs **(G, H)**. Robust unwinding is observed exclusively on substrates bearing a 3′ overhang **(I, J)**, demonstrating that *AtSUV3* requires a 3′ single-stranded extension for efficient duplex unwinding.

### Recombinantly expressed *AtSUV3* resolves transcription intermediates

To address the potential role of *AtSUV3* in resolving structures characteristic of fork or transcription collapse, we used different substrates that mimic transcription collapse. Unwinding of an arrested transcription bubble suggests that as in human mitochondria, plant SUV3 resolves transcription intermediates either via the formation of a degradosome by resolving halted transcription intermediates or R-loops (30,75,76) (**Fig. 7**). RNA helicases that resolve R-loops resulting from transcription–replication conflicts or persistent RNA:DNA hybrids are a common feature in animal nuclear and mitochondrial genomes (77–79). In human mitochondria RNase H1 is additionally required for R-loop processing and faithful initiation of mitochondrial DNA replication (80)

**Fig. 7:**
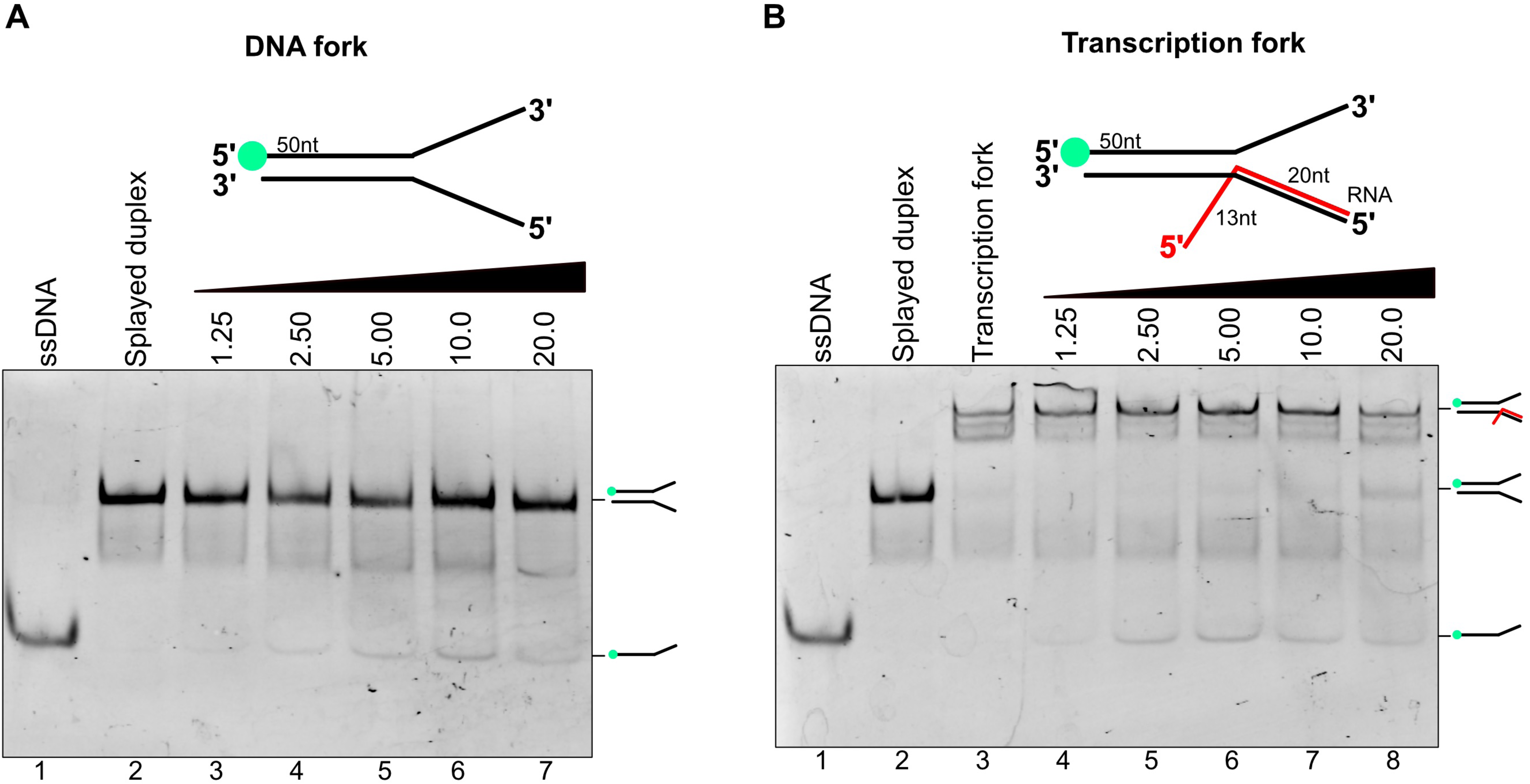
AtSUV3 unwinds forked and transcription-associated DNA structures. **A)** Helicase assay using a DNA fork substrate containing a 50-nt duplex region and non-complementary 3′ and 5′ single-stranded arms, mimicking a replication- or recombination-associated fork. Increasing concentrations of *AtSUV3* (1.25–20 µM) were incubated with the substrate and products were resolved on a native polyacrylamide gel. Strand displacement is evidenced by the appearance of the labeled single-stranded product (lanes 3 to 7). **B)** Helicase assay using a transcription fork substrate that mimics a collapsed or arrested transcription bubble. The substrate contains a 50-nt DNA duplex and a 20-nt RNA annealed to one strand, generating a 13-nt RNA 3′ overhang reminiscent of an R-loop or stalled transcription intermediate. Increasing concentrations of *AtSUV3* (1.25–20 µM) promote unwinding of the RNA–DNA hybrid and release of the labeled strand (lanes 4 to 8).

## DISCUSSION

We initiated this study to find proteins required for double stranded break repair in *Arabidopsis* mitochondria. Our screen found mutants hypersensitive to ciprofloxacin, which is a common way of inducing double-strand breaks in mitochondria and asses effects on the phenotype of mutants, or the effects of increased breakage on the dynamics of the genome (29,63,68,81). Through this screen, *AtSUV3* was identified as playing a role in DNA maintenance in plant mitochondria. *Suv3* (<u>su</u>ppressor of <u>v</u>ar1-<u>3</u>) was first discovered in *S. cerevisiae* as a dominant suppressor allele that caused overaccumulation of group I and omega introns and encodes an ATP-dependent RNA helicase (31,32).

### AtSUV3 is involved in mitochondrial DNA repair

Herein we isolated three different alleles of *SUV3*; each present slightly different phenotypes. The first allele, *suv3^K147^*, was the result of an EMS screen corresponding to a G-C to A-T transition mutation causing a missense mutation from a glycine to an arginine at position 252. This is the most C-terminal of the three alleles, and although it appears to disrupt function, it has an incompletely penetrant, semidominant phenotype. Because of the complex phenotype and consequent difficult genotyping (presumably due to the production of a misfolded protein), we sought a loss of function allele by first looking at a T-DNA insertional mutation. This insertion is located slightly upstream of the missense allele, right at the boundary between the two RecA domains (**Fig. 1**). It is not clear how stable the N-terminal portion of the protein might be, or if it interacts with other monomers, however we were unable to isolate homozygotes of this allele. The failure to recover a stable homozygous *AtSUV3* T-DNA insertion line may be due to the specific location of disruption, potential residual activity, or insertion-associated effects, any of which could influence developmental outcomes. To further investigate *AtSUV3* function genetically, we designed guide RNAs to generate a CRISPR/Cas9-induced mutation that prematurely truncates the protein near the beginning of the RecA1 domain **(Fig. 2).** This truncation ends with 8 changed amino acids and is missing so much of the protein that it is unlikely to have dominant negative effects. The phenotype is much less severe than *suv3^K147^*consistent with the interpretation that it is a true null mutation. Unlike the missense and the T-DNA mutations, homozygotes are viable and fertile, although the copy number of mitochondrial DNA is roughly half that of wild type (**Table 1**). Growth on soil, and germination rates were unaffected, even in the presence of ciprofloxacin, suggesting that the sensitivity of the *suv3^K147^* missense allele (which allowed us to find it in our screen) is allele specific. Root growth assays are generally seen as a sensitive readout of mitochondrial function (82). The *suv3^-^*mutants had unchanged root growth, even on ciprofloxacin, and unchanged silique lengths, again indicating a mild phenotype of the CRISPR/Cas9 induced null mutation. Plants lacking Suv3 grow normally, in contrast to the results from human cells and yeast. However, when plants were stressed, the stark difference between wild type plants and *suv3*^-^ mutants became clear. When stressed, plant mitochondrial genomes accumulate a higher amount of reactive oxygen species (83), and an accumulation of reactive oxygen species increases DNA damage and therefore increases the frequency of DSB repair. Since impairment is seen under stress in *suv3^-^*mutants, it is likely that the Suv3 protein is playing a role in alleviating RNA:DNA damage that arises from the increase in ROS accumulation. In fact, *SUV3* was shown to have increased transcription under salt stress conditions in rice (65). A common theme in plant mitochondrial genome dynamics is redundancy of proteins (27,84). With this in mind, it is plausible that mutation of Suv3 in plants does not produce effects as severe as those observed in other eukaryotes because it may function redundantly with other helicases in plant mitochondria. It may be that the lack of Suv3 only becomes obvious under stress because it is a stress-specific helicase, or because another helicase is more efficient at degrading RNA:DNA hybrids in non-stressed conditions. It is possible that AtSuv3 may be performing redundant roles with AtRecG, which has been shown to resolve RNA:DNA hybrids in *E. coli* (McGlynn and Lloyd 2000; Vincent et al. 1996).

Although *de novo* assembly of the Nanopore reads was consistent with the reported structure of *Arabidopsis* Col-0 mitochondria (85), minimap2 alignments to the reference genome showed elevated structural variants. This suggests that in the absence of SUV3 function, there is a significant, but low and probably variable level of aberrant junctions present. To examine this further we looked specifically for elevated ectopic recombination at a 249 bp repeat, known as Repeat L, that has been previously shown to be an indicator of perturbation of either the DNA maintenance and repair pathways or induction of high levels of double-strand breaks (10,15,66). We found that there was a significant increase in recombination at Repeat L in the *suv3*^-^ mutants, indicating that in normal conditions, *AtSUV3* is playing a role in preventing ectopic recombination of the genome. To verify that the biochemical activity was consistent with the prediction from the sequence, we purified a recombinant SUV3 protein from *E. coli* and performed helicase assays to define the enzyme specificity. The three major findings were that AtSUV3 is an ATP-dependent helicase, it has a strict 3’→5’ specificity, and it has a much higher activity on R-loop substrates than on DNA. Therefore, consistent with homologs in other species and its annotation, it is an RNA helicase that unwinds 3’-ends of RNA in R-loops.

In yeast, mutation of SUV3 results in a petite phenotype associated with loss of mitochondrial function and/or mtDNA (forming petite colonies) (57), supporting a connection between R-loop removal and mitochondrial genome maintenance. What are the possible sources of R-loops that SUV3 could act on? These could include replication primers that did not get elongated, aborted transcripts, or intermediates in retrotransposon insertion. It should be noted that Suv3 will not unwind transcription bubbles unless the 3’-end of the RNA is available, but transcription stalled at a DNA lesion or break could be a substrate for Suv3 unwinding. Another possibility is replication fork collisions with transcription bubbles, leaving structures that need to be resolved. R-loops could increase DNA damage and inhibit repair because the looped-out DNA strand is more susceptible to oxidation or deamination, the looped-out DNA strand could invade another DNA molecule as a D-loop, or the R-loops might interfere with DNA repair protein recruitment and assembly. Persisting R-loops could also cause aberrantly primed replication. Another helicase (RecG) prevents runaway replication of small circular subgenomic molecules in *Arabidopsis* mitochondria (Wallet, Le Ret et al. 2015). Suv3 may function to prevent aberrant priming by R-loops as well. It is also possible that the role of Suv3 is to remove transcription bubbles from the path of an oncoming replication fork. There is also evidence that some introns have been lost from plant mitochondrial genes, possibly by retrotranspositions (86). If the presence of an intron is important, then it is possible that Suv3 is involved in preventing reverse transcription of a mature RNA into the gene. Ultimately, the discovery that an RNA helicase is important in DNA maintenance processes is a surprising one. We note that according to the public data in ePlant, SUV3 is expressed to some extent in every tissue, but is expressed at much higher levels in stamens and pollen (87). This expression pattern could suggest a redundant gene that functions in vegetative tissue and megasporogenesis, or it could be that the specific growth and physiology of pollen development requires Suv3 to a much greater extent. We did not observe any gross fertility defects in our CRISPR-induced mutant of *SUV3*, but because pollen is generally in excess, the effect may be unobservable by just allowing plants to self-cross. Given that mitochondrial DNA in pollen is an evolutionary dead end, it is surprising that a DNA maintenance protein would be expressed there. However, it may be that removal of R-loops from the DNA is important in achieving proper DNA compaction in generative cells and degradation in post-fertilization embryos to block paternal inheritance. The *SUV3* gene merits further study for its roles in RNA maintenance, DNA maintenance and pollen development.

### AtSUV3 is a paralog of the putative Arabidopsis SUV3 degradosome helicase

The unexpected physiological role and biochemical properties of AtSUV3 raise the question of whether this protein also functions as a component of the plant mitochondrial RNA degradosome. In fungi and animals, SUV3 is a mitochondrial ATP-dependent 3′→5′ RNA helicase that serves as a core constituent of this complex, cooperating with mitochondrial exoribonucleases to promote the degradation of structured RNA substrates. In yeast and *Candida albicans*, SUV3 forms a heterodimer with the 3′→5′ exoribonuclease Dss1, whereas in humans it associates with polynucleotide phosphorylase (PNPase) to assemble the mitochondrial degradosome (55). In addition to AtSUV3 (At4g14790), which we identify here as a novel factor involved in mitochondrial DNA repair and maintenance, *Arabidopsis thaliana* encodes a second SUV3-related protein, AT5G39840.1, which we refer to here as SUV3-like. At SUV3-like protein harbors an extended N-terminal in which the first 160 amino acids after the predicted mitochondrial targeting sequence are not present in AtSUV3. The presence of this extended N-terminal region strongly resembles human and yeast SUV3. AtSUV3 and SUV3-like are highly similar proteins, sharing 39% amino acid identity and 75% similarity within their aligned regions, suggesting that they originated from a common ancestral helicase and may have evolved distinct functions (**Fig. 4S**). Phylogenetic analysis separated plant SUV3 paralogs into two distinct clades, corresponding to SUV3 and SUV3-like proteins. The presence of SUV3-like proteins in green algae suggests that the gene duplication event giving rise to these paralogs occurred early in the evolution of the green lineage, before the emergence of land plants. The retention of both protein groups throughout plant evolution implies that SUV3 and SUV3-like diversified to acquire distinct functions (**Fig. 8A**).

**Fig. 8.**
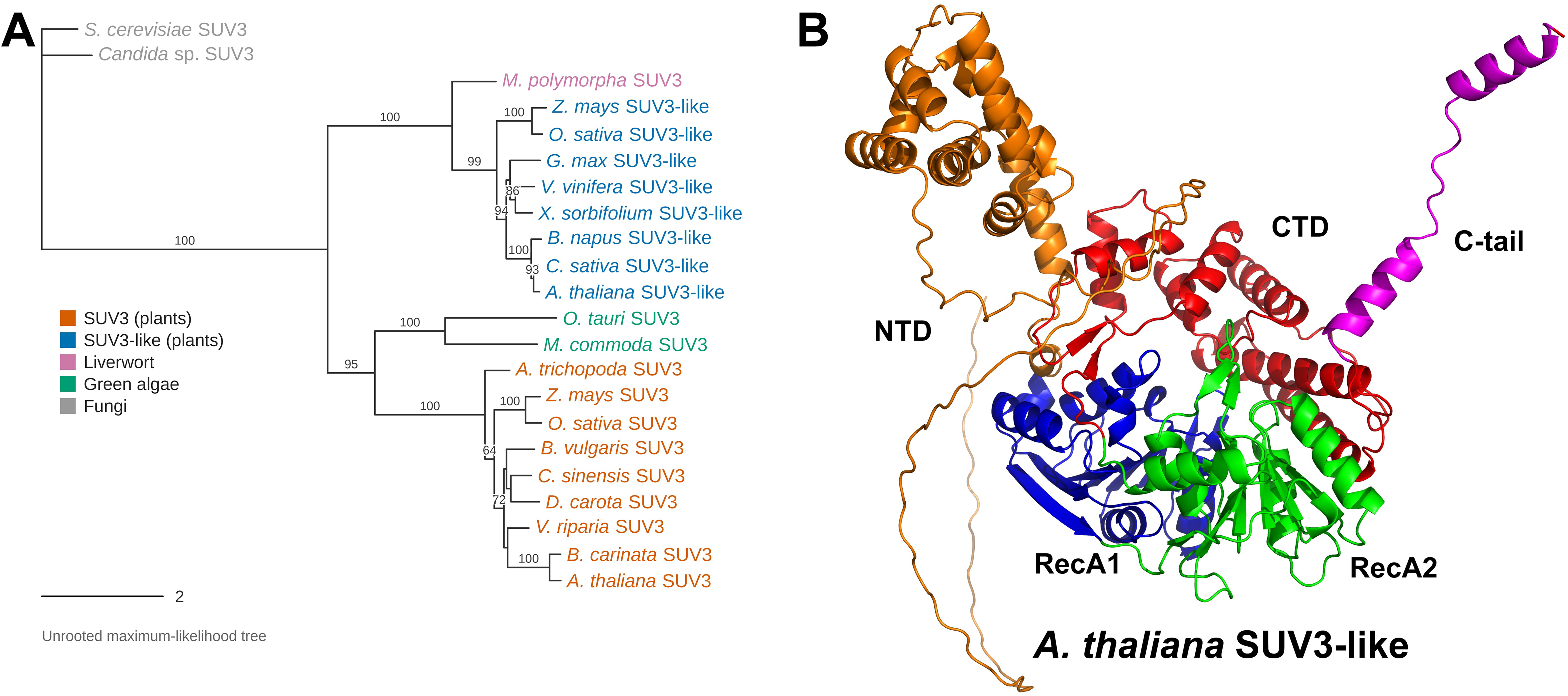
Evolutionary and structural analyses of Arabidopsis SUV3 paralogs. **A)** Unrooted maximum-likelihood phylogenetic tree showing the separation of plant SUV3 proteins into two distinct groups, corresponding to SUV3 and SUV3-like proteins. The SUV3-like group includes sequences from the liverwort *Marchantia polymorpha* and diverse angiosperms, whereas the analyzed green algal sequences cluster with canonical SUV3 proteins, consistent with an ancient duplication followed by divergence of the two plant paralogs. **B)** AlphaFold-based structural model of *Arabidopsis thaliana* SUV3-like. The conserved RecA1 and RecA2 helicase domains are shown in blue and green, respectively, with the N-terminal domain (NTD) and C-terminal domain (CTD) indicated. The SUV3-like protein also contains an extended C-terminal tail (C-tail; magenta), highlighting structural differences from canonical AtSUV3

**Fig. S3.**
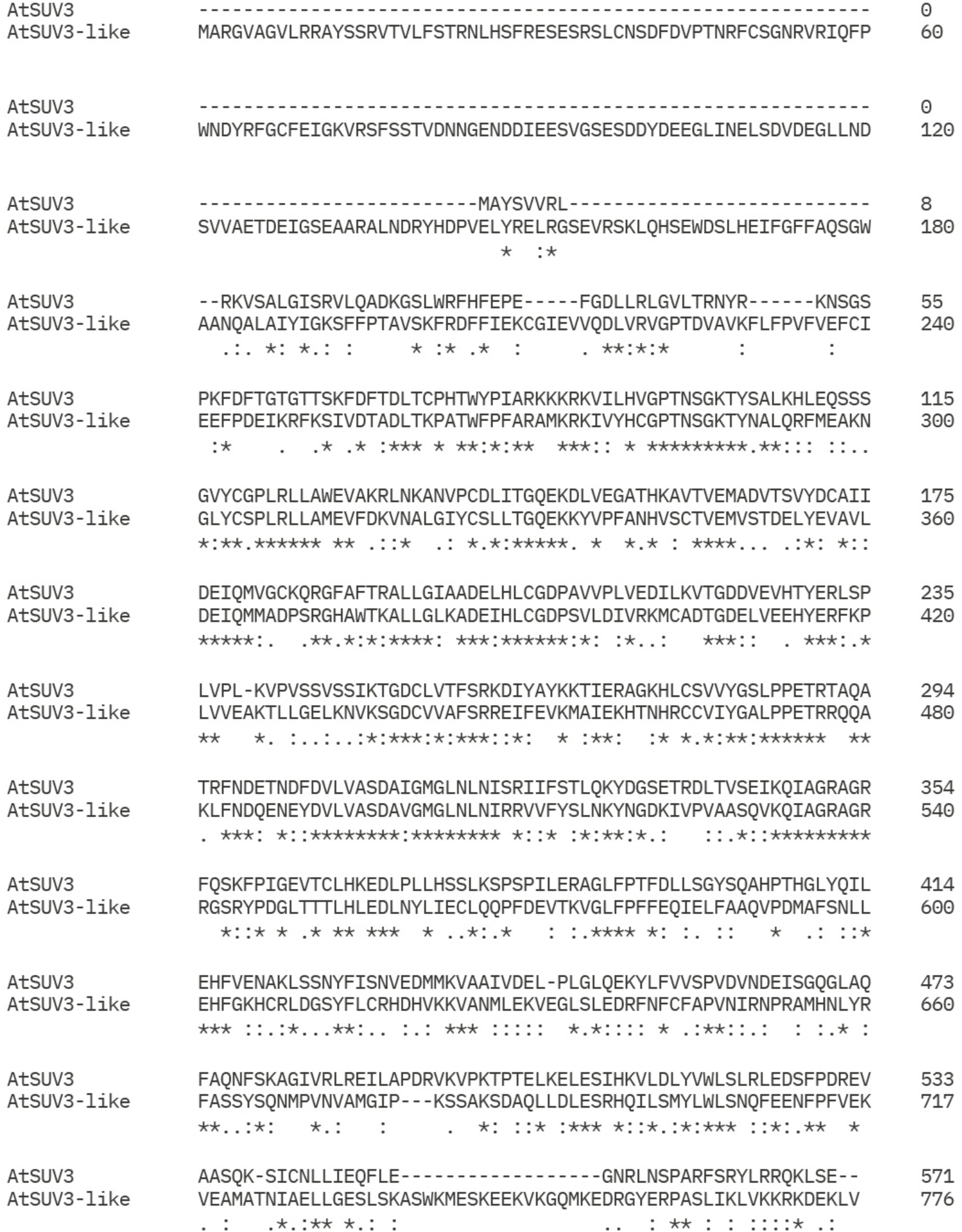
Sequence alignment of AtSUV3 and SUV3-like proteins from *Arabidopsis thaliana*. SUV3-like (AT5G39840.1) contains an extended N-terminal region of approximately 160 amino acids absent from AtSUV3 but resembling those of yeast and human SUV3 proteins.

Paralogs are abundant in plant organelles. Examples include the organellar DNA polymerases POLIA and POLIB, the single-stranded DNA-binding proteins SSB1 and SSB2, the recombinases RECA1, RECA2, and RECA3, and the four organellar single-stranded DNA-binding proteins OSB1–OSB4 (4,27,88). Gene duplication events following specialization may permit the origin of new enzymatic activities, for instance DNA polymerase POLIB now specializes in lesion bypass (4,89). Our results raise the possibility that the two SUV3 paralogs in plant mitochondria evolved new activities suggesting a model in which SUV3-like, retained the ancestral role as the helicase component of the plant mitochondrial degradosome, while AtSUV3 has undergone functional specialization toward mitochondrial DNA repair. Supporting the existence of a plant degradosome, *Arabidopsis* encodes a Dss1-like exoribonuclease, AtmtRNaseII (At5g02250), which efficiently degrades linear RNA substrates but is impeded by structured RNAs, implying a requirement for an associated helicase (90). AtSUV3-like preserves the characteristic domain organization of SUV3 proteins, including a mitochondrial targeting sequence, an N-terminal domain, two RecA-like helicase domains, and a C-terminal domain. Collectively, these findings support an evolutionary scenario in which duplication of an ancestral plant *SUV3* gene was followed by subfunctionalization, resulting in two specialized mitochondrial helicases: one involved in RNA turnover and degradosome activity, and the other, AtSUV3, functioning in the repair and maintenance of the mitochondrial genome. To our knowledge, this represents the first example of an RNA helicase family traditionally associated with RNA surveillance being implicated in mitochondrial DNA repair in plants after specialization.

## ACKNOWLEDGEMENTS

We thank Corina Díaz-Quezada for help in protein purification. We are grateful to Michaela Bonnett, Emily Kahlandt and Jake Kingsley for assistance with the cipro screening of EMS mutants, to Anne Britt for advice on mutant screens and for discussions of DNA repair in plants during ACC’s sabbatical, to Anne Britt and Mily Ron for advice on CRISPR and vectors, and to Beth Rowan, Isabelle Henry and Emily Wynn for advice on mapping by sequencing and help with Python scripting.

## AUTHOR CONTRIBUTIONS

Moira G. Rodriguez: Investigation, Writing, Validation, Formal analysis, Methodology Claudia D. Raygoza: Investigation, Validation, Formal analysis, Methodology

Ana M. Martinez-Hottovy: Investigation, Validation, Formal analysis, Methodology David M. Campbell: Investigation, Validation, Formal analysis, Methodology Emma R. Purfeerst: Investigation, Validation, Formal analysis, Methodology

Luis G. Brieba: Writing—review & editing, Funding

Alan C. Christensen: Investigation, Conceptualization, Writing—review & editing, Funding, Validation, Formal Analysis, Methodology

## DATA AVAILABILITY

Raw DNA sequencing reads for *suv3*^-^CRISPR mutant whole genome sequencing with Nanopore and Illumina,wild-type Nanopore reads, and suv3^K147^ Illumina are available at NCBI BioProject ID: PRJNA1482635 (ID 1482635 - BioProject - NCBI). Raw data for *suv3*^-^ CRISPR mutant root length measurements, silique length measurements, germination rate measurement, and repeat recombination qPCR are in Additional File 1.

---

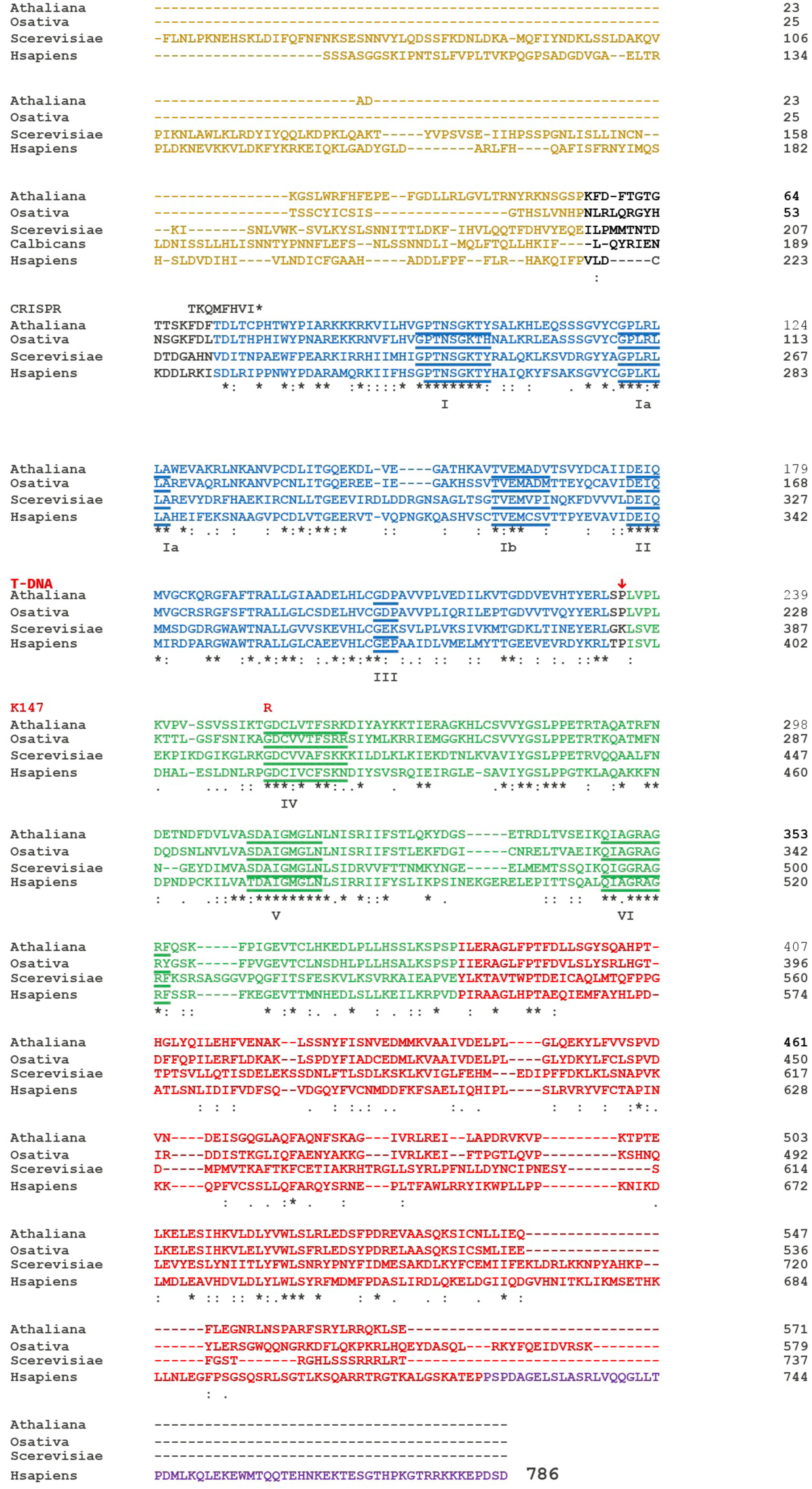

