## Supplemental information for "A specialized SUV3 RNA helicase is involved in plant mitochondrial DNA repair"

**SUPPLEMENTARY FILES**

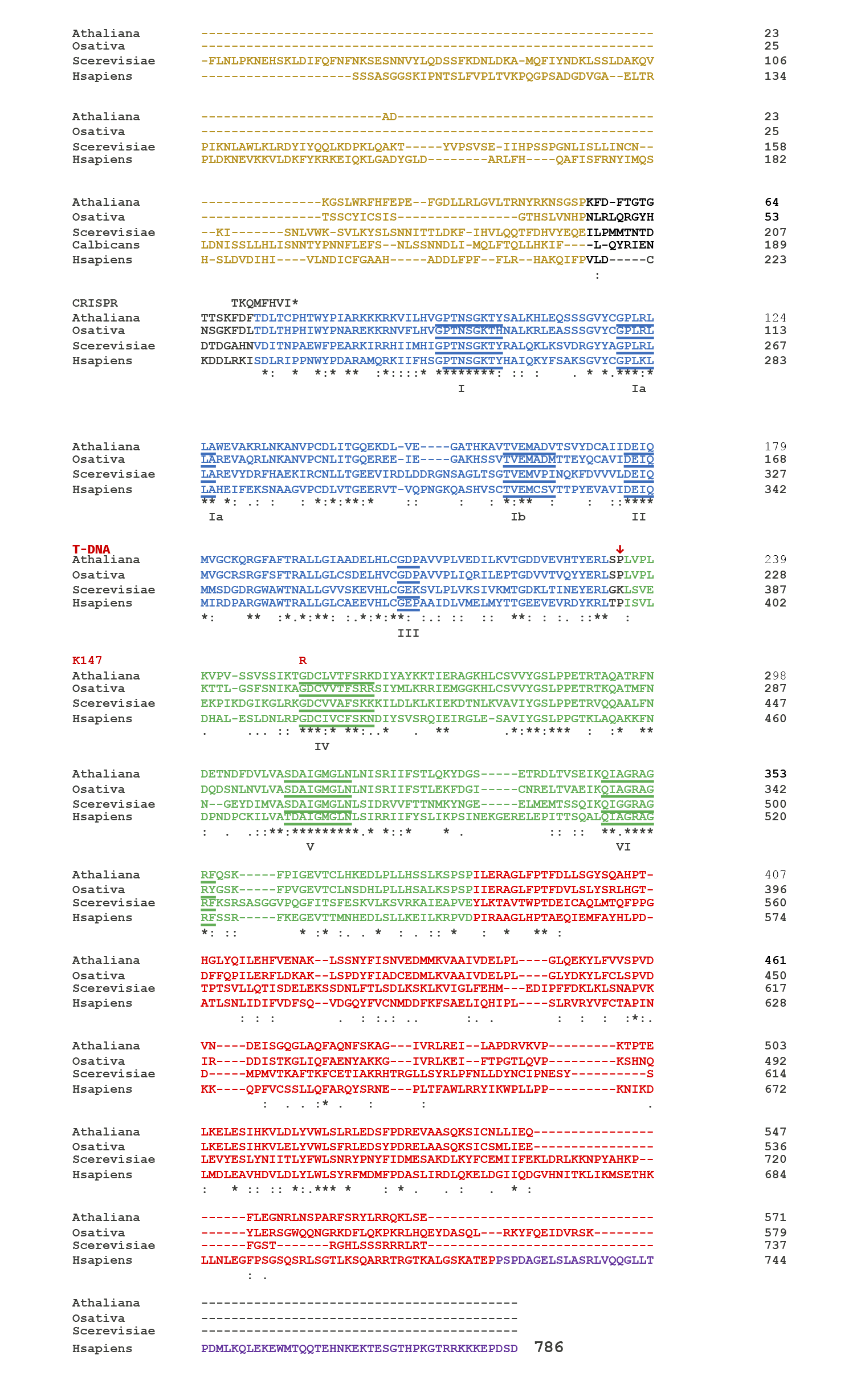

**Fig. 1S:** Amino acid sequence alignment of SUV3 orthologs from plant, human, and yeast. Conserved RecA-like helicase motifs characteristic of Superfamily 2 (SF2) helicases are indicated, and structural domains are annotated based on the crystal structure of human SUV3. Domain organization is color-coded as follows: the N-terminal domain (NTD) in gold, RecA1 in blue, RecA2 in green, the C-terminal domain (CTD) in red, and the unique C-terminal extension present in human SUV3 in purple. The mitochondrial targeting sequences are absent in this alignment (Jedrzejczak, Wang et al. 2011). Changes in the K147, T-DNA and CRISPR alleles are indicated above the *A. thaliana* sequence.

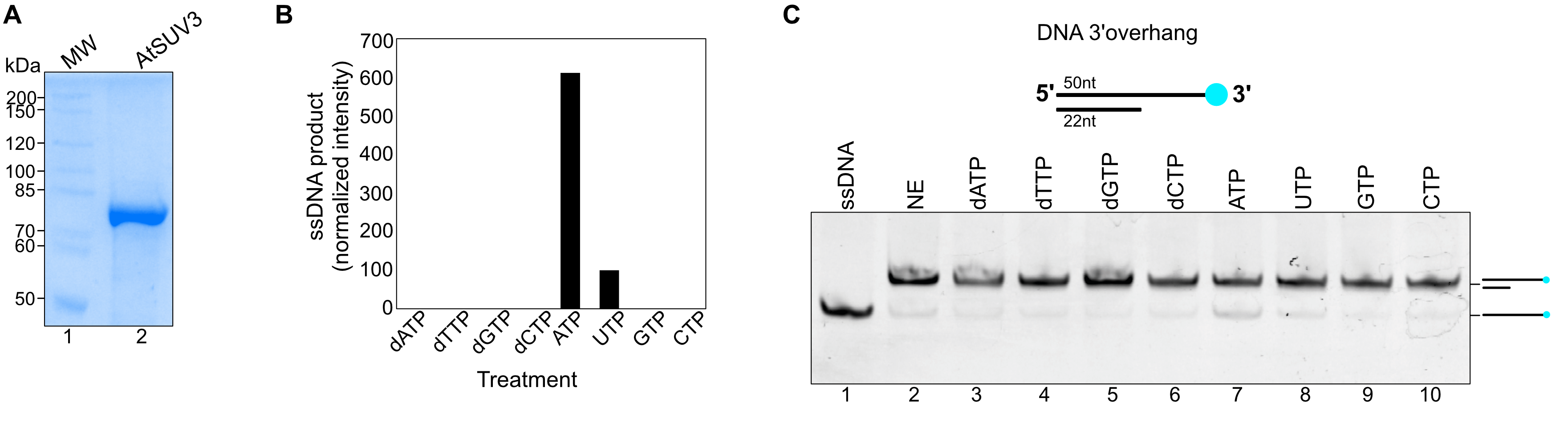

**Fig. 2S AtSUV3 is an ATP dependent 3′** → **5′ helicase A)** 10% SDS–PAGE showing the purified recombinantly expressed *AtSUV3* protein stained with Coomassie Blue. **B)** Helicase activity by *AtSUV3* requires ATP. Quantification of helicase activity represented as a bar graph depicts duplex unwinding under different nucleotides. Robust unwinding is observed only in the presence of ATP, whereas no detectable activity is measured in reactions lacking other nucleotides. **C)** Representative native gel showing the unwinding by AtSUV3 on a 3´overhang in comparison to a control with no enzyme (NE) (lane 2)

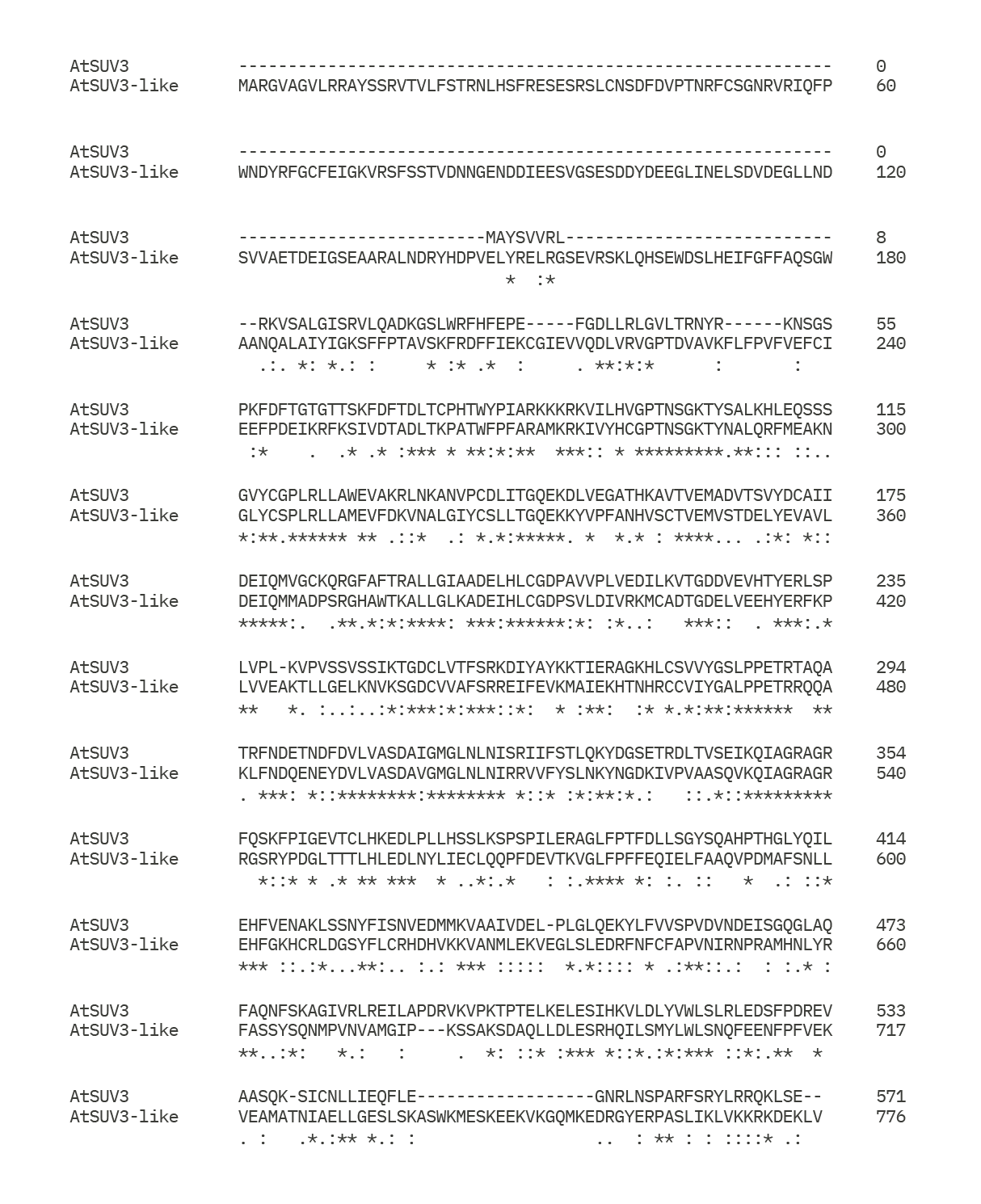

**Fig. S3 Sequence alignment of AtSUV3 and SUV3-like proteins from *Arabidopsis thaliana*.** SUV3-like (AT5G39840.1) contains an extended N-terminal region of approximately 160 amino acids absent from AtSUV3 but resembling those of yeast and human SUV3 proteins.

**Table S1. Oligonucleotides used for AtSUV3 cloning and biochemical substrate preparation**

| Name | Sequence |
| --- | --- |
| SUV3_FW | 5’-GGCATGCCATATGCTGCAAGCGGATAAGGGGAGC-3’ |
| SUV3_RV | 5’-GATTGGCGGATCCCTACTCACTCAATTTCTG-3’ |
| PHO | Cy5-5’-ATAAATATTTTTTATTAATAATAGATCACCTTTCTTTCTCTTCTCCCCTT-3’ |
| PHO_COMP | 5’-AAGGGGAGAAGAGAAAGAAAGGTGATCTATTATTAATAAAAAATATTTAT-3’ |
| OH-5OH | 5’-ATTATTAATAAAAAATATTTAT-3’ |
| RT_COMPL | Cy5-5’-GATGTTAGCCATGACCTA-3’ |
| SUV3_3OVER | Cy5-5’-TAGCTGAGTCCAGTCTTA-3’ |
| 3_OVERH_RD | 5’-TAAGACTGGACTCAGCTAGGTCATGGCTAACATC-3’ |
| 3_OVERH_R | 5’-UAAGACUGGACUCAGCUAGGUCAUGGCUAACAUC-3’ |
| ARM_D | FAM-5’-CCGCTACCAGTGATCACAGGTCATGGCTAATGCTGTTCATGAACCTGTTT-3’ |
| DLF | 5’-GGGTGAACCTGCAGGTGATGTTAGCCATGACCTGTGATCACTGGTAGCGG-3’ |
| ARM_R | 5’-GAACUGCAAUCGGCAUCACCUGCAGGUUCACCC-3’ |

**Table S2.** Primers used in FPNI and qPCR reactions.

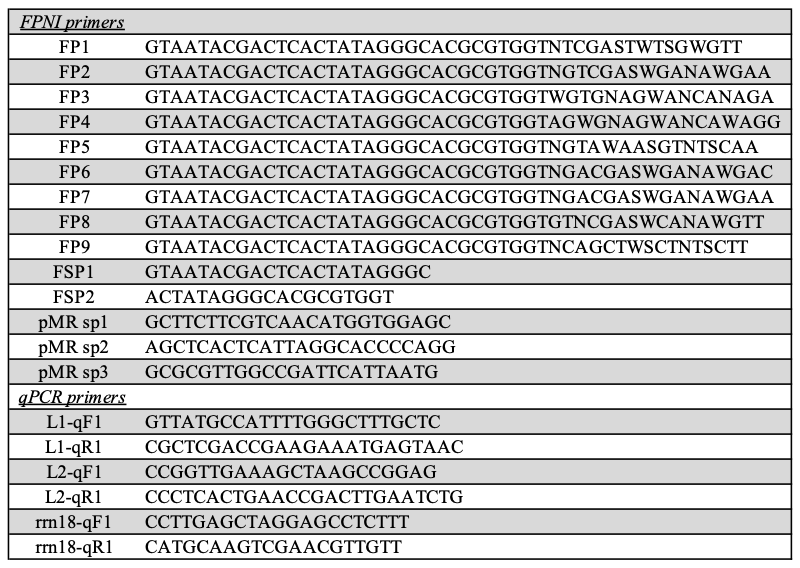
